# microRNA-22 displaces ITAFs from the 5’UTR and inhibit the translation of Coxsackievirus B3 RNA

**DOI:** 10.1101/2023.07.29.551118

**Authors:** Priya Rani, Biju George, V Sabarishree, Somarghya Biswas, Raju S Rajmani, Apala Pal, Saumitra Das

**Author notes:** Address for correspondence: Dr. Saumitra Das, Professor, Department of Microbiology and Cell Biology, Indian Institute of Science, Bangalore-560012, India.

## Abstract

microRNAs play an essential role in gene regulation during virus infections and have major consequences on viral pathogenesis. During RNA virus infections, the host miRNAs can target both host mRNAs and the virus genomic RNA. Using the CVB3 virus as a model, we have investigated how a host miRNA can target viral genomic RNA and act as an antiviral factor limiting the growth of the virus. CVB3 is an RNA virus whose infection causes myocarditis and, eventually, dilated cardiomyopathy. We shortlisted miRNAs with a potential binding site in the CVB3 genomic RNA. Among these, miR-22 was picked for further studies as its binding site was putatively located in a region in the CVB3 5’ UTR, important for recruiting ITAFs and ribosomes for IRES-mediated translation. Using mutational analysis and pull-down assays, we first confirmed the binding of miR-22 on the 5’UTR. This binding negatively regulated the translation of CVB3 RNA. However, miR-22 binding-defective mutant of CVB3 RNA had no effect of miR-22 overexpression and could translate normally. Moreover, cells from which miR-22 was knocked out, showed a higher level of CVB3 infection as compared to the wild type. We have further demonstrated that the binding of miR-22 interferes with the recruitment of several ITAFs (La, PSF, and PTB) on viral mRNA. This abrogates the spatial structure necessary for ribosome recruitment on the CVB3 RNA, ultimately inhibiting its translation. Also, the level of miR-22 increases 4 hours post-infection, presumably after the synthesis of viral 2A protease, to regulate infection in the host cell more effectively. Along with the direct effect on viral RNA, the altered level of miR-22 affects the level of its cellular targets which might contribute to CVB3 infection. To identify the possible players, we obtained a list of miR-22 targets and performed pathway analysis. Several targets were shortlisted among the top hits and their levels upon CVB3 infection were checked. Protocadherin-1 (PCDH-1), a single-pass transmembrane protein, followed an expected trend, and its levels were significantly downregulated upon CVB3 infection in miR-22 dependent manner. miR-22 mediated suppression of PCDH1 levels during CVB3 infection points towards the possible role of miR-22 in either modulating antiviral signaling or in virus entry, in addition to regulating the viral translation.

## INTRODUCTION

Coxsackievirus B3 (CVB3) is a positive-strand RNA virus from the *Picornaviridae* family that spreads through the fecal-oral route. It is a human pathogen infecting the heart, pancreas, and brain causing myocarditis, acute pancreatitis, and aseptic meningitis, respectively. The inflammation caused by viral infection leads to long-term fatal diseases such as dilated cardiomyopathy and type-I diabetes. To establish infection, CVB3 is completely dependent on the host machinery, and the host factors such as proteins and non-coding RNAs play a major role in it.[1-4]

Among the non-coding RNAs, miRNAs have been implicated in influencing CVB3 life cycle by many ways. miRNAs can exert both direct and indirect effects during viral infections[5]. The indirect mechanism involves their control over viral replication by targeting specific genes within the host genome. In CVB3 infection, miRNAs play critical roles in regulating viral replication and host responses. Upregulated miRNAs like miR-203 enhance viral replication and cell survival by targeting ZFP-148[6]. Additionally, miR-590-5p in vesicles from infected cells prolongs viral replication by targeting SPRY1[7]. miR-126 upregulation modulates the ERK1/2 and Wnt/b-catenin pathways, promoting virus replication and release[8]. Conversely, miR-221/-222 suppress target genes to limit viral replication and inflammation, protecting the heart during CVB3 infection[9]. Moreover, during CVB3 infection, the virus carefully manipulates cell apoptosis to facilitate its replication and spread. Specific miRNAs, such as miR-34a, miR-222, miR-98, and miR-21, play crucial roles in regulating this process. MiR-34a acts as a pro-apoptotic factor, while miR-222, miR-98, and miR-21 act as anti-apoptotic factors in viral myocarditis[10-14]. There are reports suggesting the direct involvement of miRNAs during CVB3 infection. Bioinformatic analysis has revealed that miR-342-5p potentially targets the 2C-coding region of the CVB3 viral genome, effectively inhibiting viral replication by directly acting on this region. Interestingly, not all miRNAs function as negative regulators[15]. Another miRNA, miR-10a∗, plays a unique role in promoting viral biosynthesis by targeting the 3D-coding region (nt6818-nt6941) of CVB3. Furthermore, miR-10a∗ is found to be abundant in the heart of BALB/c mice, suggesting its potential influence on viral myocarditis (VMC) by enhancing CVB3 replication[16]. These findings indicate the potential utility of miRNAs as a valuable treatment approach to directly impede viral replication by specifically targeting crucial regions of the viral genome.

The direct influence of miRNAs on the CVB3 life cycle piqued our interest, considering the high likelihood of viral RNA being targeted by host miRNAs. Consequently, our focus shifted towards identifying miRNAs that may exhibit binding affinity towards the untranslated regions of the viral RNA. Using the VITA database, we successfully obtained a list of miRNAs with a strong binding probability to both the 5’ and 3’ UTRs of CVB3. The corresponding list is presented in Table S1. Notably, one of the miRNAs, miR-22, had a predicted binding site within the 5’ UTR, precisely between stem loop V and VI (Figure 1A). This is a crucial region in the CVB3 5’ UTR, as it is known to facilitate interactions with several host RNA-binding proteins, subsequently promoting CVB3 RNA translation[17, 18]. Additionally, this region contains a Shine-Dalgarno-like sequence, which serves as the recruitment site for the small subunit of the ribosome to start scanning, leading to translation initiation at the initiator AUG codon[19].

**Figure 1.**
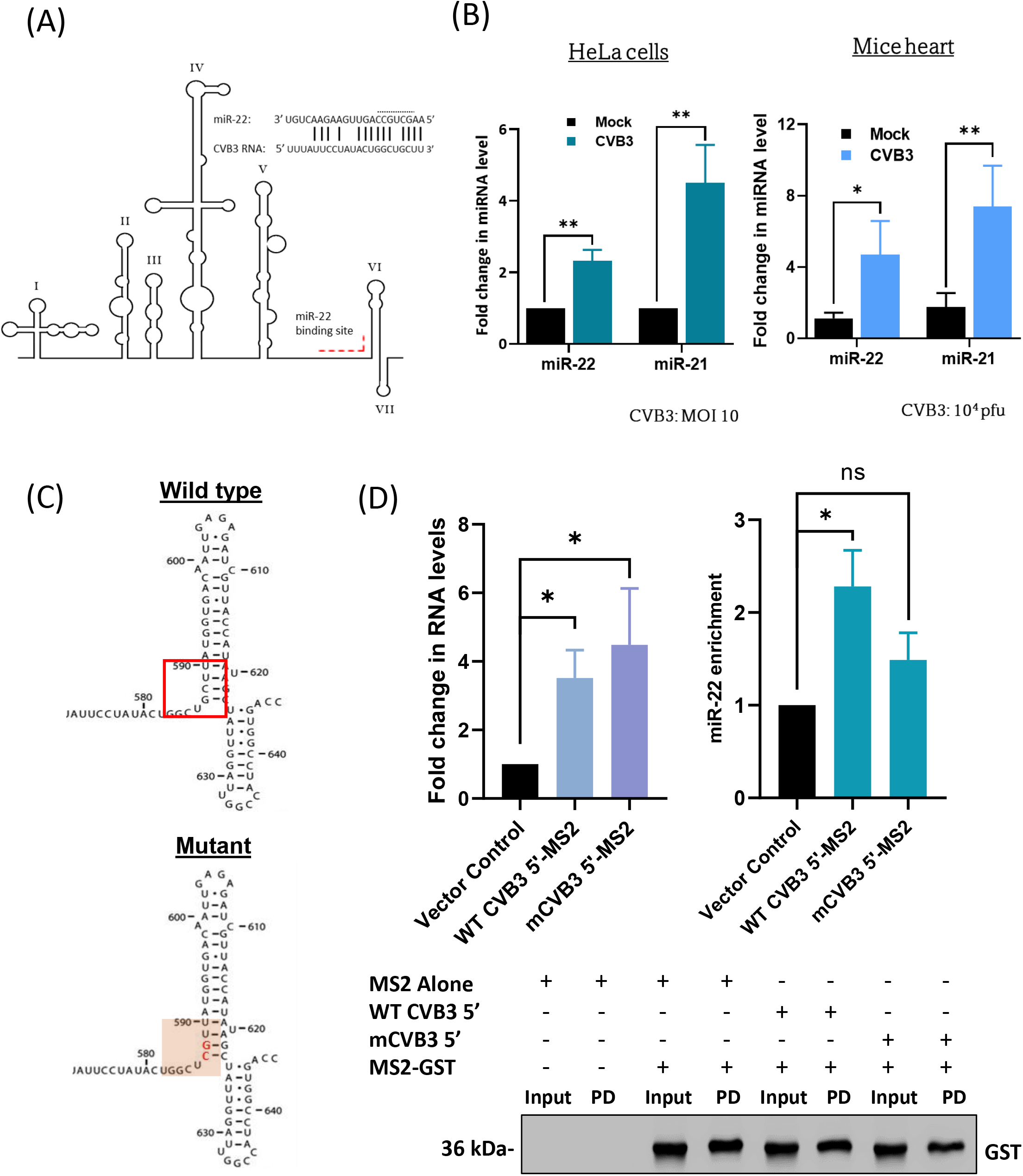
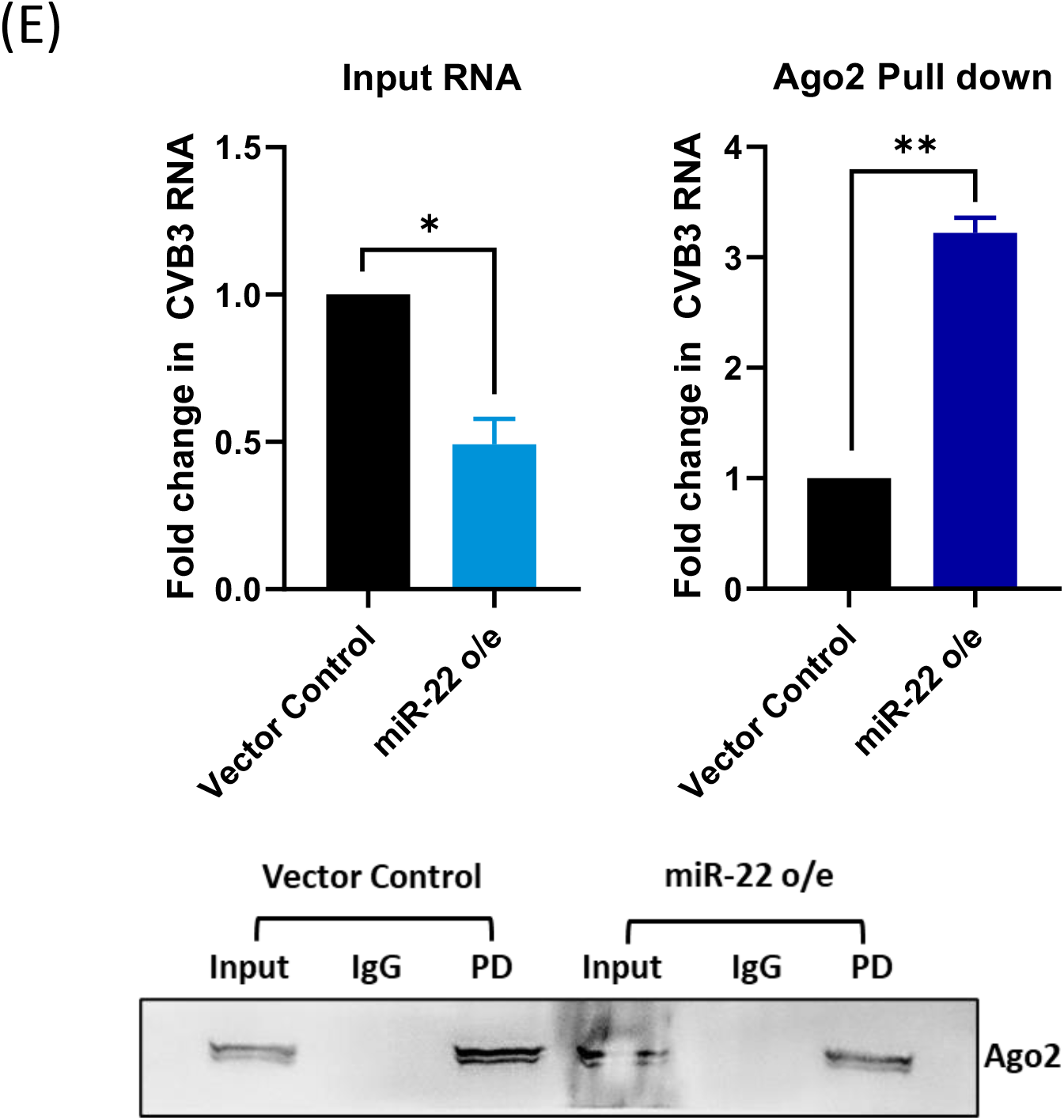
miR-22 levels are induced upon CVB3 infection, and it binds to the 5’UTR of CVB3 RNA. A) Schematic representation of predicted miR-22 binding site in the CVB3 5’UTR. B) HeLa cells were seeded at a confluency of 90% and infected with CVB3 (MOI 10). Cells were harvested 8 h.p.i. and total RNA was isolated from the lysate. miR-22 and miR-21 levels were checked using qPCR. Error bars represent standard deviation in three independent experiments. 3-4 weeks old male BALB/c mice were injected with 104 pfu of CVB3 intraperitoneally. 7 days post infection, the animals were sacrificed, and the mice hearts were harvested. Total RNA was isolated from the tissues and miR-22 and miR-21 levels were checked using qPCR. Error bars represent standard deviation in five biological replicates. C) Schematic representation of miR-22 binding mutation in the CVB3 5’UTR. The represented stemloop is sl VI. D) CVB3 5’UTR WT or miR-22 binding mutant overexpression construct and pMS2-GST constructs were co-transfected in HeLa cells. The bar graph on the left shows the level of overexpression of the CVB3 5’UTR constructs. The bar graph on the right shows the miR-22 enrichment in the pull-down fractions using qPCR. Western blot analysis was done from the input and pull-down cell extracts and probed with GST antibody. E) HeLa cells were transfected with either pSuper (vector control) or miR-22 overexpression contruct and pull down was done using Ago-2 antibody from the cellular extract harvested post 8 hours of transfection. The bar graph represents the qPCR analysis of CVB3 RNA present in the miR-22 overexpression cells as compared to the vector control. The RNA was normalized with the input CVB3 RNA present in the extract. Western blot was done to confirm the pull-down. Error bars represent standard deviation in three independent experiments. ***=p value<0.001, **=p value<0.01, *=p value<0.05.

miR-22 is a well-studied miRNA with diverse roles in various disease conditions. In the context of cancer, it acts as a tumor suppressor[20-23], while in arthritis, it exhibits anti-inflammatory properties[24]. Downregulation of miR-22 levels has been observed in Alzheimer’s disease, contributing to protein aggregate formation and disease progression[25, 26]. Of particular interest, miR-22 has also been implicated in several cardiac pathologies. It is abundantly expressed in the heart, and studies have revealed that when miR-22 is knocked out in mice hearts and cardiac stress is given to the mice, it results in the development of cardiac dilation and fibrosis, a phenotype similar to the pathogenesis induced by CVB3 infection. Moreover, miR-22 levels are increased during cardiac hypertrophy, highlighting its crucial role in the cardiac remodeling process[27, 28].

Given the intriguing properties of miR-22, we further explored its potential role during CVB3 infection.

## RESULTS

### miR-22 levels are induced upon CVB3 infection, and it binds to the 5’UTR of CVB3 RNA

First of all, we were interested to check the level of miR-22 upon CVB3 infection, as a miRNA that was upregulated had a higher chance of binding to CVB3 RNA and regulate its functions. We used two model systems for checking miR-22 levels post CVB3 infection. HeLa cells, a well-established cell culture model to study CVB3 life cycle and BALB/c mice (3-4 weeks old) as an *in vivo* model system. qPCR was done to check miR-22 levels from the total RNA isolated from CVB3-infected HeLa cells and mice heart tissues. We found that in the case of both HeLa cells as well as mice heart tissues, miR-22 levels were upregulated upon CVB3 infection. miR-21 is used in our study as a positive control (Figure 1B).

Since miR-22 was upregulated during CVB3 infection and also had a predicted binding site in the CVB3 5’UTR, we first wanted to confirm that miR-22 was actually binding to the CVB3 RNA. In order to do so, we generated miR-22 binding mutant (Figure 1C) in the CVB3 5’UTR-MS2 RNA (TRAP construct) and performed TRAP assay to check the presence of miR-22 upon pull-down. As can be clearly observed in Figure 1D, the pull-down fraction of WT but not miR-22 binding mutant CVB3 5’UTR showed enrichment of miR-22. We also confirmed the binding by performing pull down experiment using Ago-2 antibody upon mir-22 overexpression and found that more CVB3 RNA can be detected in the pull-down upon miR-22 overexpression (Figure 1E). The results indicate that miR-22 binds to the 5’UTR of CVB3 RNA.

### miR-22 negatively regulates CVB3 life cycle

We were next interested in checking how miR-22 binding is regulating the CVB3 life cycle. To explore it, we utilized a CVB3 replicon construct, that has Firefly luciferase gene in place of the CVB3 structural genes. It is a non-infective, replicating CVB3 genome. We overexpressed or partially silenced miR-22 in HeLa cells and transfected them with CVB3 replicon RNA. Cells were harvested after 8 hours of transfection and luciferase activity was measured. We found that upon miR-22 overexpression luciferase activity of CVB3 replicon was reduced. Similarly, upon partial knockdown of miR-22, the luciferase activity increased (Figure 2B). We further validated these findings using a CVB3 replicon defective in the binding of miR-22. We observed that while miR-22 overexpression decreased the luciferase activity of WT CVB3 replicon, the mutant replicon had no effect (Figure 2C). These results suggest negative regulation of CVB3 life cycle by miR-22.

**Figure 2.**
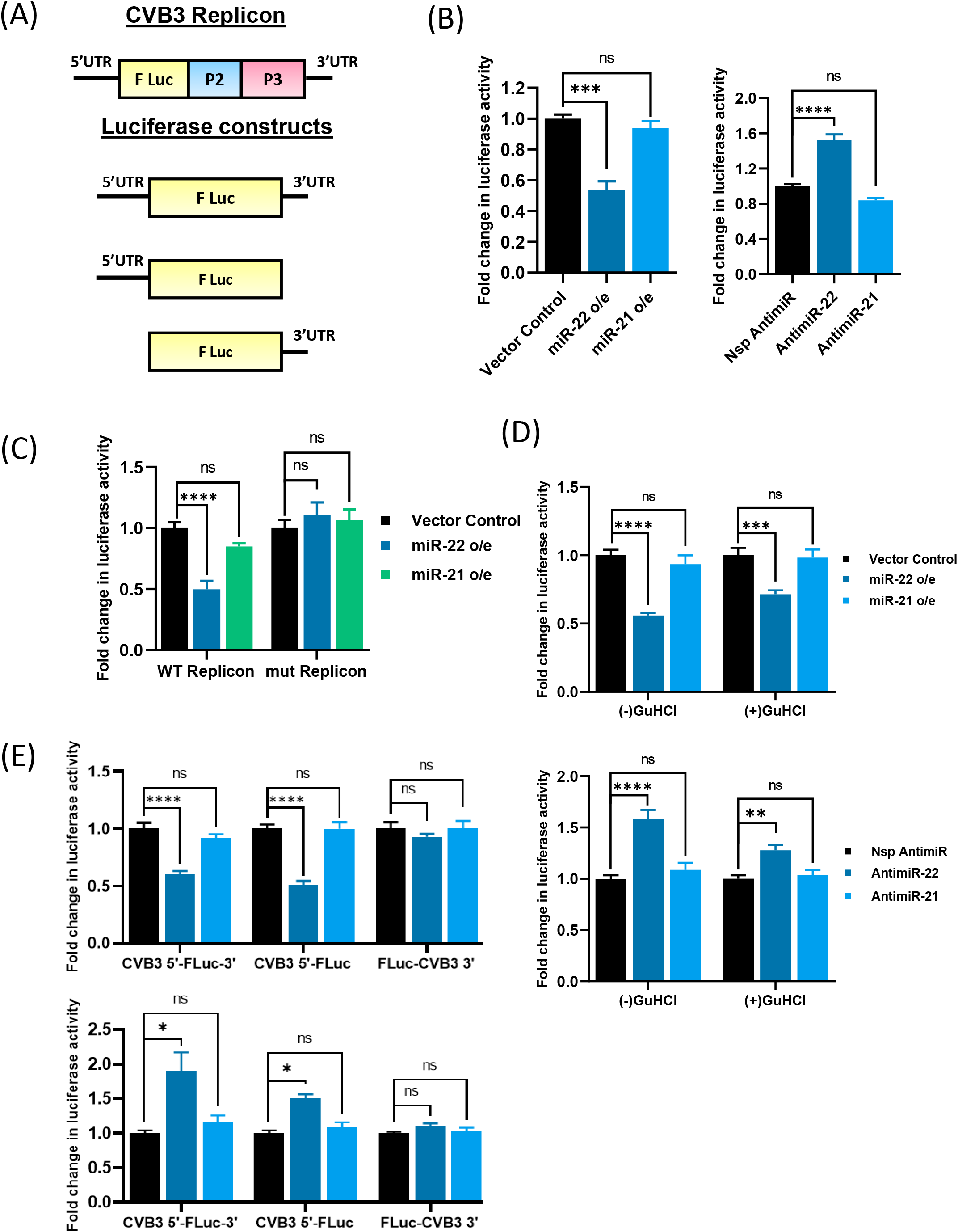
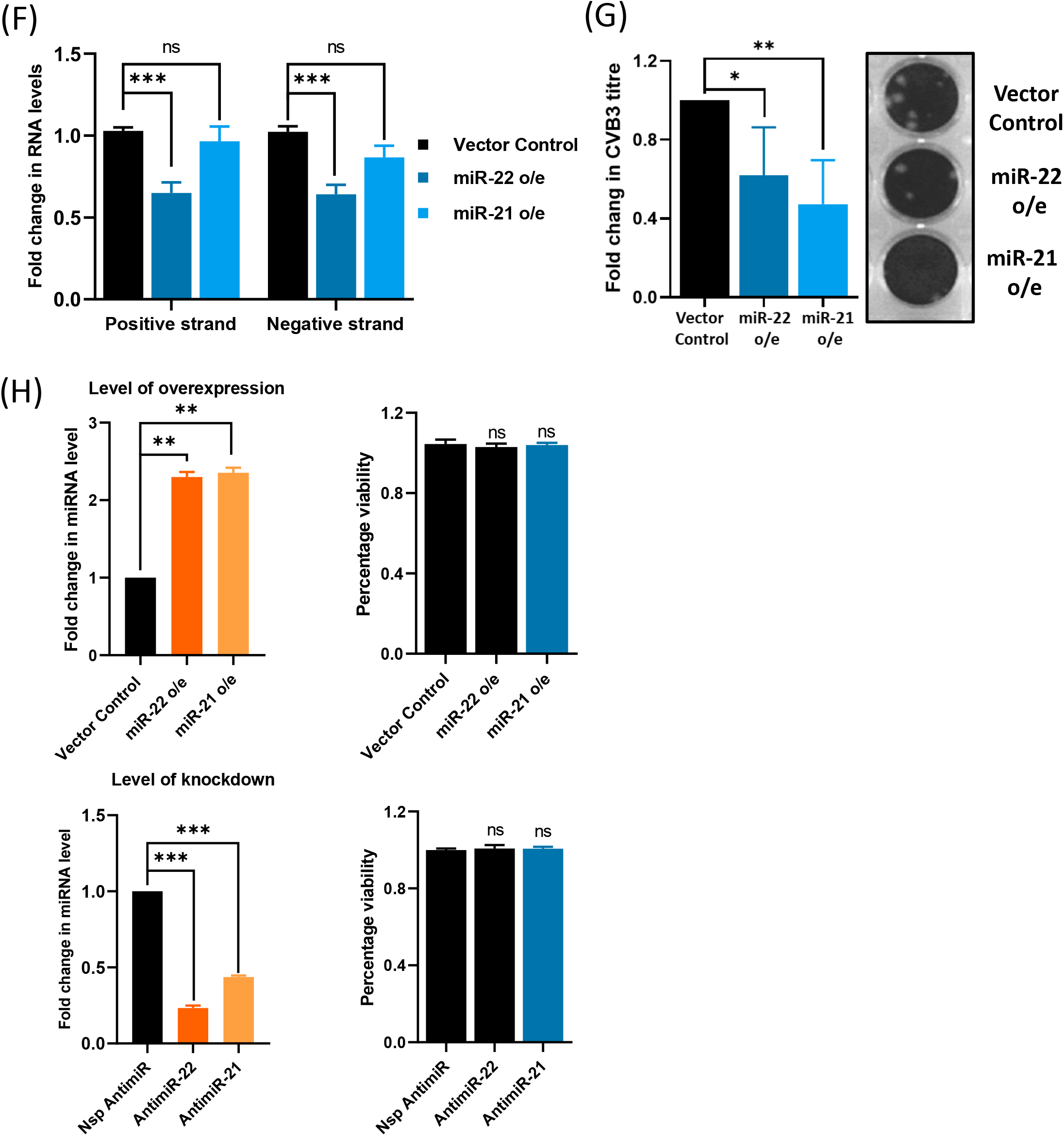
miR-22 negatively regulated CVB3 life cycle. A) Schematic representation of the luciferase constructs used in the experiments. B) HeLa cells were transfected with pSuper (Vector control), miR-22 or miR-21 overexpression plasmids for overexpression experiment and non-specific Anti-miRNA, AntimiR-22, or AntimiR-21 for partial knockdown experiment. After 24 hours, CVB3 replicon RNA was transfected, and the cells were harvested after 8 hours followed by checking of the luciferase activity. The bar graph on the left represents the fold change in luciferase activity of CVB3 replicon upon miR-22 or miR-21 overexpression. The bar graph on the right represents the fold change in Luciferase activity of CVB3 replicon upon partial knockdown of miR-22 or miR-21. C) HeLa cells with miR-22 overexpressed were transfected with either WT or miR-22 binding defective CVB3 replicon RNA and harvested after 8 hours. Fold change in luciferase activity is represented as the bar graph. D) HeLa cells overexpressing vector control, miR-22 or miR-21 were transfected with CVB3 replicon RNA either in absence or in presence of 2mM GuHCl in media. Cells were harvested 8 hours post transfection and luciferase activity was measured (Upper panel). HeLa cells transfected with Nsp Anti-miR, AntimiR-22 or AntimiR-21 were further transfected with CVB3 replicon RNA either in absence or in presence of 2mM GuHCl in media. Cells were harvested 8 hours post transfection and luciferase activity was measured (Bottom panel). E) HeLa cells overexpressing vector control, miR-22 or miR-21 were transfected with each of the described luciferase construct RNAs. Cells were harvested 8 hours post transfection and luciferase activity was measured. Bar graph represents the fold change in Luciferase activity (Upper panel). HeLa cells transfected with Nsp AntimiR, AntimiR-22 or AntimiR-21 were further transfected with each of the described luciferase construct RNAs. Cells were harvested 8 hours post transfection and luciferase activity was measured. Bar graph represents the fold change in Luciferase activity (Bottom Panel). F) HeLa cells overexpressing vector control, miR-22 or miR-21 were infected with CVB3 at MOI 10. Cells were harvested 8 hours post transfection and total RNA was isolated from the cell. CVB3 RNA was quantified by qPCR. The bar graph represents the fold change in the positive and negative strand of CVB3. G) HeLa cells overexpressing vector control, miR-22 or miR-21 were infected with CVB3 at MOI 10. Supernatant was collected post 8 hours of infection. The supernatant was used to infect a monolayer of Vero cells and after 1 hour of adsorption, layered with 1.6% low-melting agarose. After 48 hours, cells were fixed with 4% PFA, agarose was removed, and plaques were stained using Crystal violet dye and counted. The bar graph represents the fold change in the virus titre. H) HeLa cells were transfected with either vector control, miR-22 or miR-21 overexpression plasmids, or with Nsp AntimiR, AntimiR-22 or AntimiR-21 and 24 h post-transfection, MTT reagent was added to the wells, and after 3–4 h, DMEM was removed, and DMSO was added and mixed well. Reading was taken at 550 nm in a spectrophotometer. One well from each condition was kept for checking the level of overexpression of the miRNAs. The bar graphs on the left shows the level of over expression or knockdown and bar graphs on the right shows the MTT reading. Error bars represent standard deviation in three independent experiments. ****=p value<0.0001 ***=p value<0.001, **=p value<0.01, *=p value<0.05.

We were further interested in understanding at what particular step during the CVB3 life cycle, miR-22 was exerting its effect. So, we investigated the effect of miR-22 on CVB3 RNA translation by utilizing replication inhibiting drug, Guanidine Hydrochloride (GuHCl). HeLa cells overexpressing miR-22 or with partially silenced miR-22 were transfected with CVB3 replicon RNA with or without the presence of 2mM GuHCl in the media. We found that even in the presence of GuHCl, the same effect was observed on the luciferase activity of CVB3 replicon (Figure 2D). We also did another functional assay establishing the effect of miR-22 by binding specifically on CVB3 5’UTR. We used three luciferase constructs, CVB3 5’UTR-F Luc-CVB3 3’UTR, CVB3 5’UTR-F Luc and F Luc-CVB3 3’UTR. We found that miR-22 only affected the luciferase activity of the constructs having CVB3 5’UTR. These results indicated that miR-22 binding to CVB3 5’UTR had a negative effect on CVB3 RNA translation (Figure 2E).

We wanted to further confirm whether the effect we observed on CVB3 replicon and luciferase constructs reverberated with CVB3 infection as well. So, HeLa cells overexpressing miR-22, were infected with CVB3 at MOI 10 and harvested after 8 hours of infection. The cell lysate was used for total RNA isolation and the supernatant was collected for plaque assay. We found that both the positive as well as negative strand CVB3 RNA was downregulated in the cells overexpressing miR-22 (Figure 2F). Moreover, the CVB3 virus titre produced was also low (Figure 2G). MTT assay was done to confirm that the overexpression or knockdown alone had no effect on cells (Figure 2H). So, overall, it appears that miR-22 negatively regulates CVB3 at all the different steps of infection cycle.

### CVB3 proliferation is increased in miR-22 knockout cells

To further validate our results, we generated a miR-22 knockout cell line. We confirmed the knockout by sequencing as well as quantitative PCR analysis (Figure S1 and 3A). We infected WT and miR-22 KO HeLa cells seeded at 90% confluency with CVB3 at MOI 10 and harvested the cells after 8 hours. Both CVB3 positive and negative RNA strands were found to be upregulated upon infection in miR-22 KO cells as compared to WT HeLa cells, which confirms our previous results. (Figure 3B). Also, the virus titre produced from the miR-22 KO cells was approximately 3-fold higher (Figure 3C). These results indicated that miR-22 was acting as an antiviral molecule during CVB3 infection.

**Figure 3.**
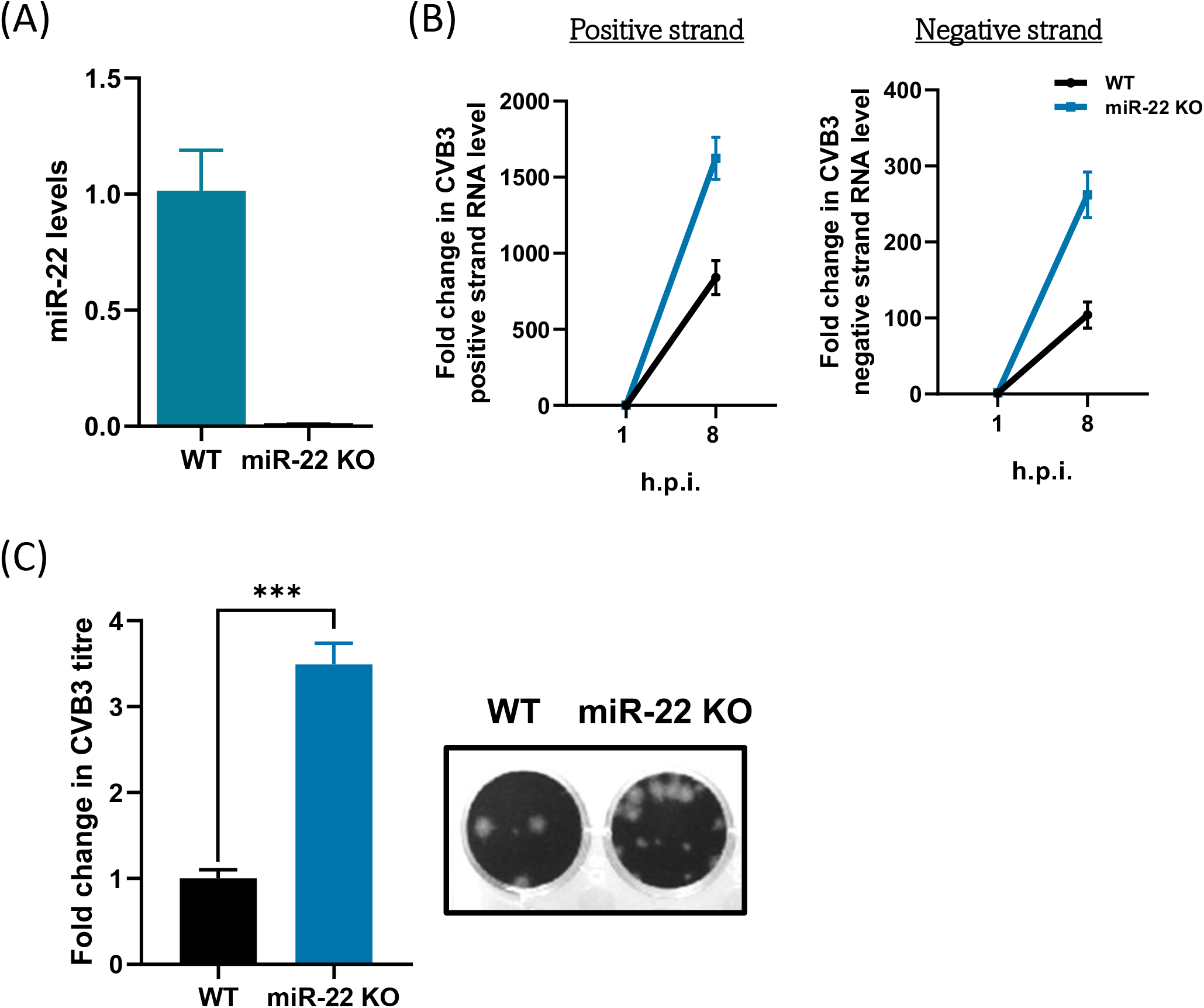
CVB3 proliferation is enhanced in miR-22 KO cells. A) Total RNA was isolated from WT HeLa and miR-22 KO HeLa cells and miR-22 levels were detected by qPCR. The bar graph represents the fold change in miR-22 levels. B) WT and miR-22 KO cells seeded at 90% confluency were infected with CVB3 (MOI 10) and harvested 1 hour and 8 hours post infection. Total RNA was isolated and CVB3 positive and negative RNA strands were quantified by qPCR. The line graphs represents the fold change in the level of CVB3 RNA in miR-22 KO cells with respect to WT cells. C) WT and miR-22 KO cells seeded at 90% confluency were infected with CVB3 (MOI 10) and the supernatant was collected 8 h.p.i. The supernatant was used to infect a monolayer of Vero cells and after 1 hour of adsorption, layered with 1.6% low-melting agarose. After 48 hours, cells were fixed with 4% PFA, agarose was removed, and plaques were stained using Crystal violet dye and counted. The bar graph represents the fold change in the virus titre. Error bars represent standard deviation in three independent experiments. ****=p value<0.0001 ***=p value<0.001, **=p value<0.01, *=p value<0.05.

### microRNA-22 binding reduces the binding of translation-promoting ITAFs

After delineating the role of miR-22 during CVB3 infection, we wanted to understand how miR-22 exerted its effect. As we discussed earlier, the site at which miR-22 binds was also a binding site of several ITAFs which promoted CVB3 translation, namely, La, PSF and PTB. We were interested to investigate how miR-22 binding affected the binding of these ITAFs to CVB3 5’UTR. So, after infecting HeLa cells overexpressing miR-22 or vector control, with CVB3, we did immunoprecipitation using antibodies against each of the ITAFs and checked the CVB3 RNA associated with them in both conditions. We also used a negative control for our experiment, HuR protein, which is another RNA binding protein harbouring a binding site in a different stem loop of CVB3 5’UTR[29]. Interestingly, we found that the CVB3 RNA associated with La, PSF and PTB was reduced upon miR-22 overexpression (Figure 4B-D). However, there was no effect on CVB3 RNA associated with HuR protein (Figure 4E). These results indicated that miR-22 was displacing the ITAFs binding to the CVB3 5’UTR between stem loop V and VI.

**Figure 4.**
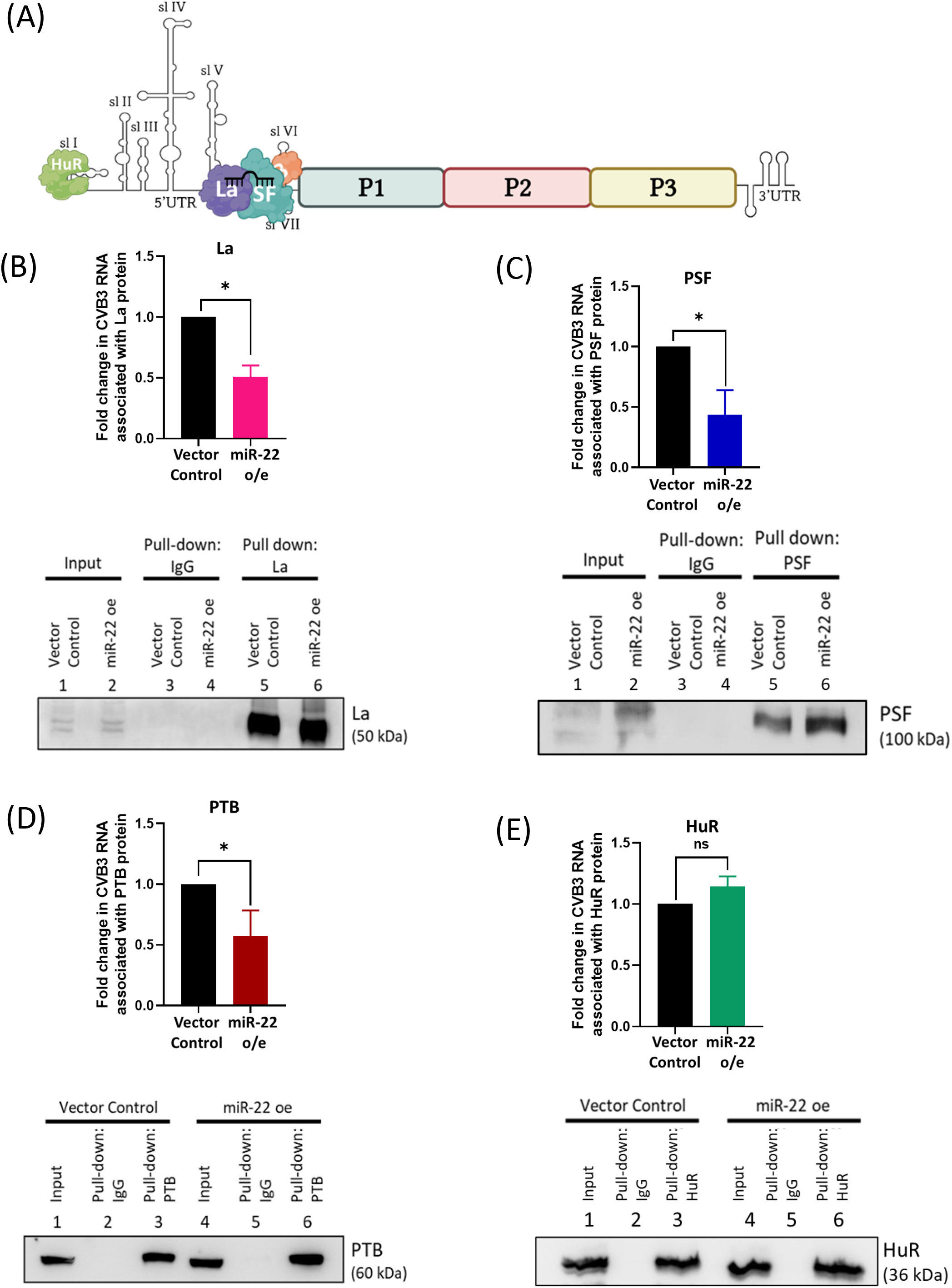
miR-22 binding to CVB3 5’UTR displaces the ITAFs. A) A pictorial representation of the binding site of miR-22(Black), La(Purple), PSF(Teal), PTB(Orange) and HuR(Green) in the CVB3 5’UTR. B, C, D and E) HeLa cells overexpressing miR-22 were infected with CVB3 (MOI 10). Cells were harvested after 8 h.p.i. and the lysate was used for immunoprecipitation using antibodies against La, PSF, PTB and HuR. Total RNA was isolated from the total lysate and the pull-down fractions and CVB3 RNA was quantified by qPCR. The bar graphs represents the CVB3 RNA associated with each of the proteins, normalized to the CVB3 RNA present in the total cell lysate. Western blots confirms immunoprecipitation. Error bars represent standard deviation in three independent experiments. ****=p value<0.0001 ***=p value<0.001, **=p value<0.01, *=p value<0.05.

### microRNA-22 binding inhibits translation by inhibiting ribosome recruitment

One of the ITAFs that were displaced by miR-22 binding to the CVB3 5’UTR was La protein. One of our previous studies has shown that La binding is essential for ribosome recruitment to the 5’UTR[19]. So, we wanted to check whether miR-22 binding was also affecting the ribosome recruitment. To explore this question, we utilised the polysome profiling technique. We checked the level of CVB3 RNA associated with the 40S ribosomal subunit, complete monosomes and polysomes, in the presence and absence of miR-22 in the cells. First, we checked whether there is any difference in the polysome profile of WT and miR-22 KO HeLa cells. As can be observed in Figure 5A, the profiles were almost identical. Next, we infected the cells with CVB3 and used the infected cell lysate to perform polysome profiling. We isolated the RNA separately from 40S fraction, the monosomes and the polysomes. As expected, we found that CVB3 RNA was present more in all the fractions (40S, monosomes and polysomes) in the case of CVB3-infected miR-22 KO cells (Figure 5C). These results indicated that miR-22 is inhibiting CVB3 RNA translation by abrogating the ribosome recruitment to the IRES region.

**Figure 5.**
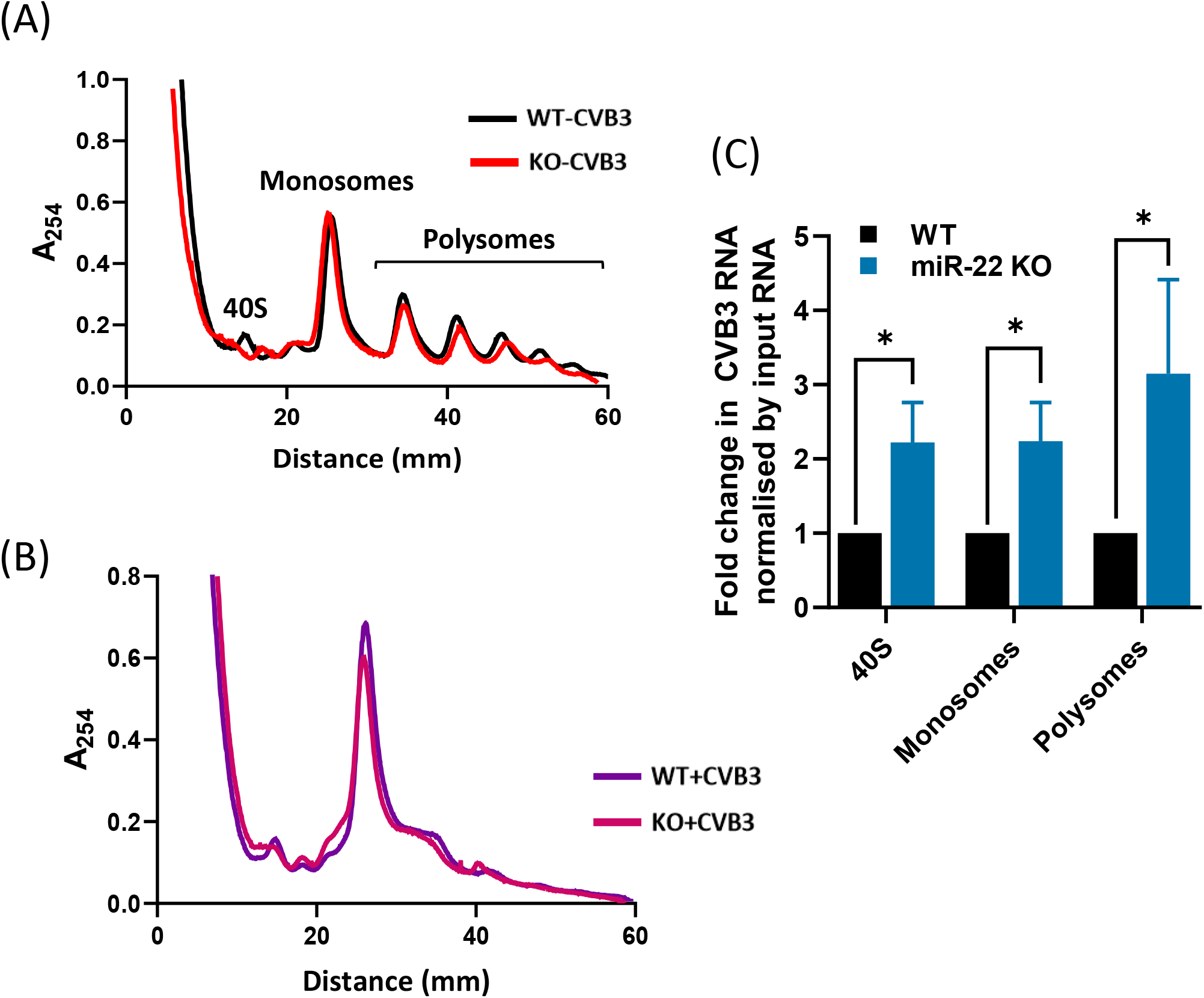
miR-22 binding to CVB3 5’UTR inhibits ribosome recruitment. A) Uninfected WT and miR-22 KO HeLa cells were used for polysome profile analysis to check global translation profile. The profiles for WT HeLa (Black) and miR-22 KO cells (Red) are shown. The 40S, Monosomes and polysomes peaks are indicated. B) CVB3 -infected WT and miR-22 KO HeLa cells were used for polysome profile analysis to check global translation profile. The profiles for CVB3 infected WT HeLa (Purple) and miR-22 KO cells (Pink) are shown. C) Total RNA was isolated from the 40S, monosomes and polysome fractions of the CVB3-infected WT and miR-22 KO cells and CVB3 RNA was quantified in them by qPCR. The bar graph represents the fold change in CVB3 positive strand present in the individual fractions in miR-22 KO cells with respect to WT HeLa cells. Error bars represent standard deviation in three independent experiments. ****=p value<0.0001 ***=p value<0.001, **=p value<0.01, *=p value<0.05.

### microRNA-22 levels increase post 4 hours of infection triggered by 2A protease

The translation of the genomic RNA strand is one of the most crucial steps in the life cycle of CVB3. As CVB3 virion particle does not carry any viral protein packaged along with the genome, it undergoes translation as soon as it enters the cytoplasm of the host cell to synthesize viral proteins. Moreover, in HeLa cells, during the first two hours of CVB3 infection, there is predominantly CVB3 RNA translation going on in the cells and a robust RNA replication starts only after 4 hours (Figure 6A). These viral proteins are utilized by the virus to replicate its genome. However, if miR-22 levels increase and do not allow CVB3 RNA to translate, it will hinder a successful infection cycle. To address this question, we checked the level of miR-22 at different time points post CVB3 infection in HeLa cells. We infected HeLa cells with CVB3 (MOI 10) and harvested the cells at 2, 4, 6, and 8 h.p.i. Interestingly, we found that the miR-22 levels increased only after 4 hours of infection allowing enough time to synthesize the viral proteins (Figure 6C).

**Figure 6.**
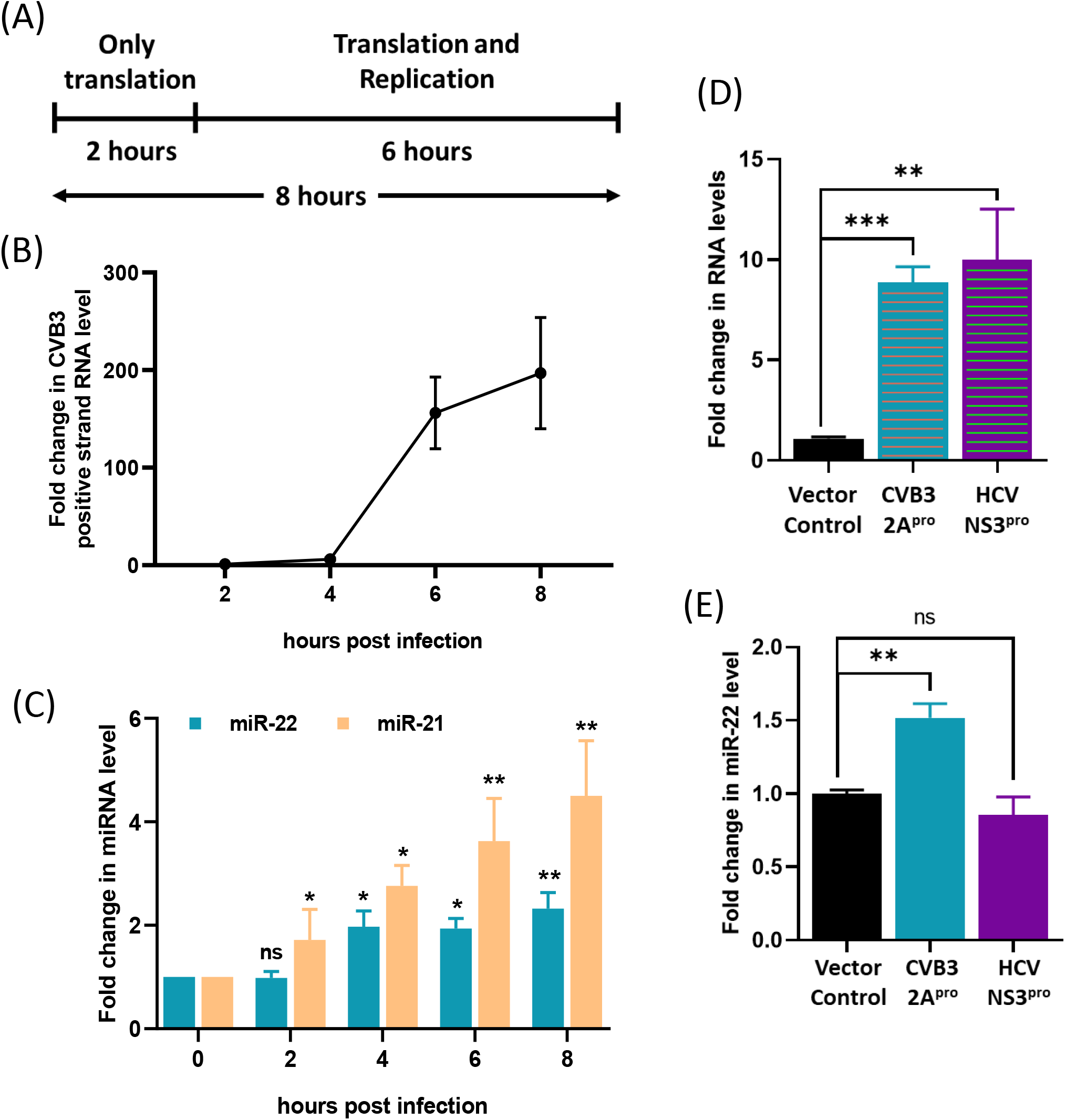
miR-22 levels increase post 4 hours of CVB3 infection as a result of 2A^pro^ accumulation. A) Schematic representation of the time course of CVB3 infection in HeLa cells. For the first two hours, there is majorly CVB3 RNA translation and after that replication and translation both picks up a robust speed. B) HeLa cells seeded at a confluency of 90% were infected with CVB3(MOI 10) and the cells were harvested post 2-,4-,6- and 8- hours of infection. Total RNA was isolated and CVB3 positive strand RNA was quantified using qPCR. C) The level of miR-22 and miR-21(positive control) were detected at the different time points post CVB3 infection. The blue bar represents the level of miR-22 and the orange bar represents the level of miR-21. D) HeLa cells seeded at a confluency of 60-70% were transfected with either pcDNA3 (Vector Control), CVB3 2Apro or HCV NS3pro expression plasmids and 24 hours post transfection, cells were harvested and total RNA isolated. The RNA level of CVB3 2Apro or HCV NS3pro were quantified using qPCR. E) The RNA level of miR-22 was quantified using qPCR. Error bars represent standard deviation in three independent experiments. ****=p value<0.0001 ***=p value<0.001, **=p value<0.01, *=p value<0.05.

Around 4 h.p.i., viral proteins start to accumulate in the cell cytoplasm. One of them being CVB3 protease, 2A. 2A protease is known to be involved in regulating many host proteins by either cleaving them or modulating their levels. Recently, one of our studies showed that it downregulated the level of a miRNA, miR-125b-5p, during CVB3 infection[29]. So, we were curious to check whether 2A^pro^ accumulation had anything to do with the increased level of miR-22 during CVB3 infection. We expressed 2A protease in HeLa cells and checked the level of miR-22. We also used NS3 protease from HCV as a control. We found that 2A^pro^ expression in HeLa cells leads to an increase in miR-22 levels (Figure 6E). These results indicated that miR-22 levels during CVB3 infection are triggered by 2A protease.

### Indirect effect of miR-22 on CVB3 infection: Protocadherin-1 is regulated by miR-22 during CVB3 infection

Since the upregulated levels of miR-22 during CVB3 infection will also regulate the host mRNAs that are targeted by miR-22, we wanted to investigate the indirect effect of miR-22 on CVB3 infection through host mRNA targets. We got a list of 825 targets of miR-22 from miRbase database and did pathway analysis of the targets using Panther software (Figure S2). From the pathway analysis, we picked the top three pathways and shortlisted the genes common between them. Subsequently, we had a list of 12 mRNAs which were possibly targeted by miR-22. Table S2 shows the list of those targets.

We first checked the mRNA levels of the shortlisted miR-22 targets upon CVB3 infection (Figure 7A). Since miR-22 is upregulated upon CVB3 infection, we only shortlisted those genes which were downregulated for further investigation. Among the 12 genes, we found four that showed inverse correlation, namely, Protocadherin-1 (PCDH1), Guanine nucleotide-binding protein subunit beta-4 (GNB4), TGF-beta receptor type-1 (TGFBR1), and Protein Wnt-5a (WNT5A). Next, we wanted to check their level in miR-22 KO cells. If the genes are regulated by miR-22, their levels should be more in miR-22 KO cell line. Interestingly, we found the levels of PCDH1 and GNB4 were upregulated in miR-22 KO cell line. Further, upon CVB3 infection in miR-22 KO cells, we found that GNB4 was showing the same trend that it was showing during infection in WT cells. However, PCDH1 was showing the opposite trend (Figure 7B). We also checked the level of PCDH1 in CVB3-infected mice heart and we found that similar to HeLa, PCDH1 levels were downregulated (Figure 7C). These results suggested that PCDH1 might be regulated by miR-22 during CVB3 infection.

**Figure 7.**
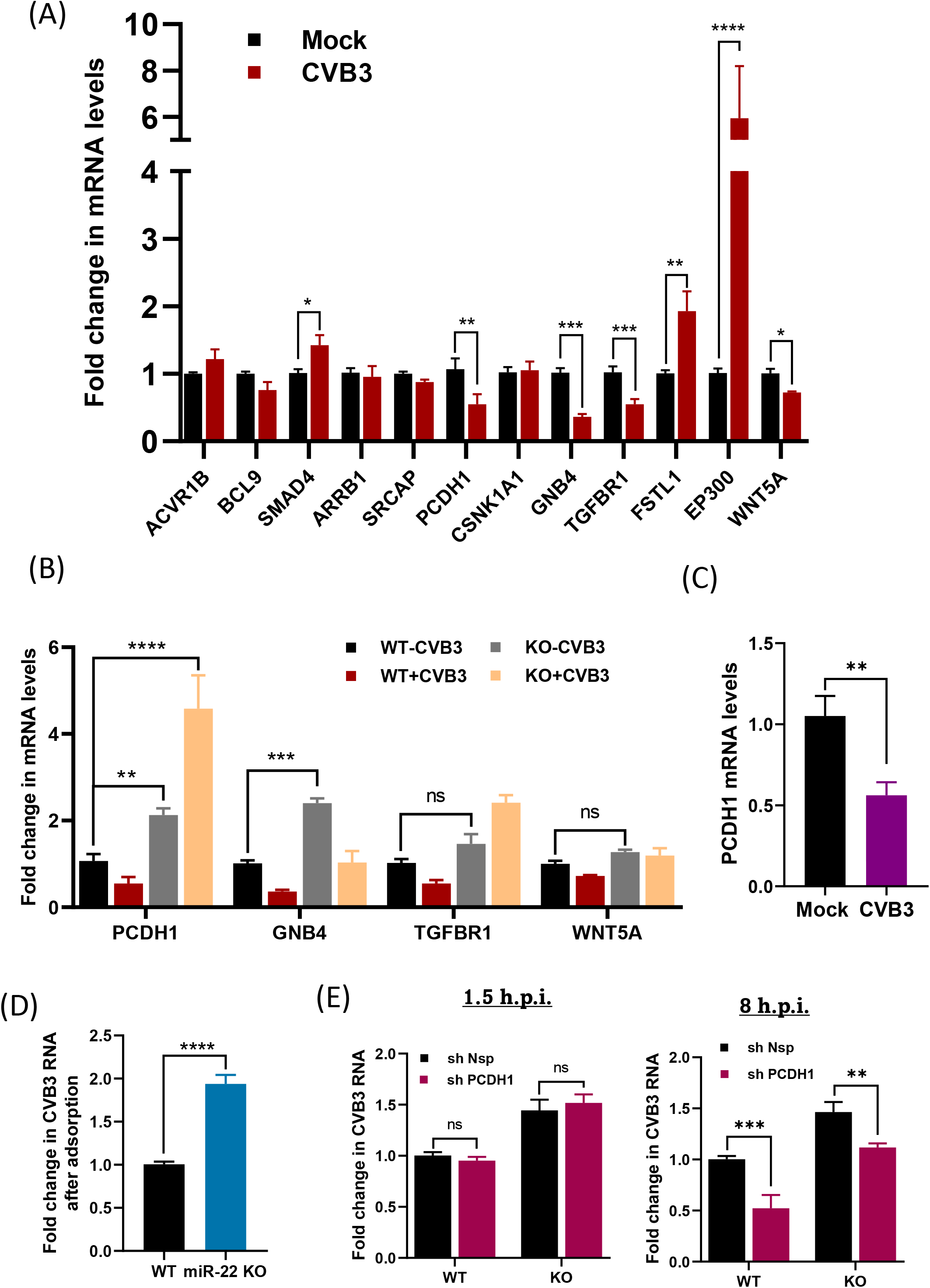
Protocadherin-1 is regulated by miR-22 during CVB3 infection and acts as a pro-viral factor. A) HeLa cells seeded at a confluency of 90% were infected with CVB3(MOI 10) and the cells were harvested post 8 hours of infection. Total RNA was isolated and the mRNA levels of different miR-22 target genes shortlisted in table 3.2 were quantified by qPCR. The bar graph represents the level of mRNA targets in CVB3 infected cells v/s uninfected cells. B) WT and miR-22 KO HeLa cells were seeded at a confluency of 90% and were either mock-infected or CVB3-infected (MOI 10). Cells were harvested after 8 h.p.i. and total RNA was isolated. mRNA levels of PCDH1, GNB4, TGFBR1 and WNT5 were quantified by qPCR. C) PCDH1 mRNA levels were quantified in CVB3-infected mice heart with respect to mock-infected mice heart post 7 days of infection. D)WT and miR-22 KO cells seeded at 90% confluency were infected with CVB3 (MOI 10) and harvested after 1 hour of adsorption. Total RNA was isolated, and CVB3 positive strand RNA levels were quantified by qPCR. The bar graph represents the fold change in CVB3 that entered in miR-22 KO cells with respect to WT cells. E) WT and miR-22 KO cells seeded at 50% confluency were transfected by either sh Nsp or sh PCDH1 plasmids and after 24 hours, infected with CVB3 (MOI 10). Cells were harvested 1.5- and 8-h.p.i. and total RNA isolated. CVB3 RNA was quantified in the cells by qPCR. Error bars represent standard deviation in three independent experiments. ****=p value<0.0001 ***=p value<0.001, **=p value<0.01, *=p value<0.05.

### Protocadherin-1 promotes CVB3 infection

Protocadherin-1 (PCDH1) is a fairly underexplored protein. It is a single-pass transmembrane protein present in cell-cell junctions. There are two interesting reports about PCDH1. One report showed that it activates NF-κB signaling in case of pancreatic cancer[30]. Another report showed that PCDH1 helps in the entry of the New World hantaviruses into the host cells[31]. Based on these reports we hypothesized that during CVB3 infection, when the virion encounters a host cell, PCDH1 present in the vicinity might get activated and in turn activated some signaling or it might act as a co-receptor and help CVB3 in its entry. To our surprise, we did find that CVB3 entry was more in miR-22 KO cells as compared to WT HeLa cells (Figure 7D). So, we partially silenced PCDH1 in miR-22 KO cells and checked its effect on CVB3 infection. We observed that while at an early time phase, there was no effect of PCDH1 knockdown on CVB3 infection, at a later time during infection, it downregulated the CVB3 infection (Figure 7E). These results indicated that PCDH1 promoted CVB3 infection.

### microRNA-22 levels post-infection determine the virus replication in various organs

Now that we established the direct and indirect roles of miR-22 during CVB3 infection, we also wanted to check if it had any role in deciding the tissue/organ tropism during CVB3 infection. miR-22 is differentially expressed in different organs of the body, being highly expressed in the heart. Moreover, different cell types have different cellular milieu that decides their response to a viral infection. So, upon infection, different organs might show differential regulation of miR-22. This might ultimately decide how CVB3 proliferates in different organs. To perform this experiment, we infected 3-4 weeks old male BALB/c mice with 10^4^ pfu CVB3 or only PBS (Mock) and sacrificed the animals after 7 d.p.i. We harvested different organs and confirmed the CVB3-induced pathogenesis by doing histopathology of heart tissue. We found fibrotic tissue and enlarged nuclei indicating cardiac injury in CVB3-infected mice (Figure 8B). Next, we checked the expression of miR-22 in the different organs. Interestingly, we found that miR-22 was downregulated post infection only in small intestine, which is also the primary target organ of CVB3, however in rest of the organs, miR-22 levels were either upregulated or unchanged (Figure 8D).

**Figure 8.**
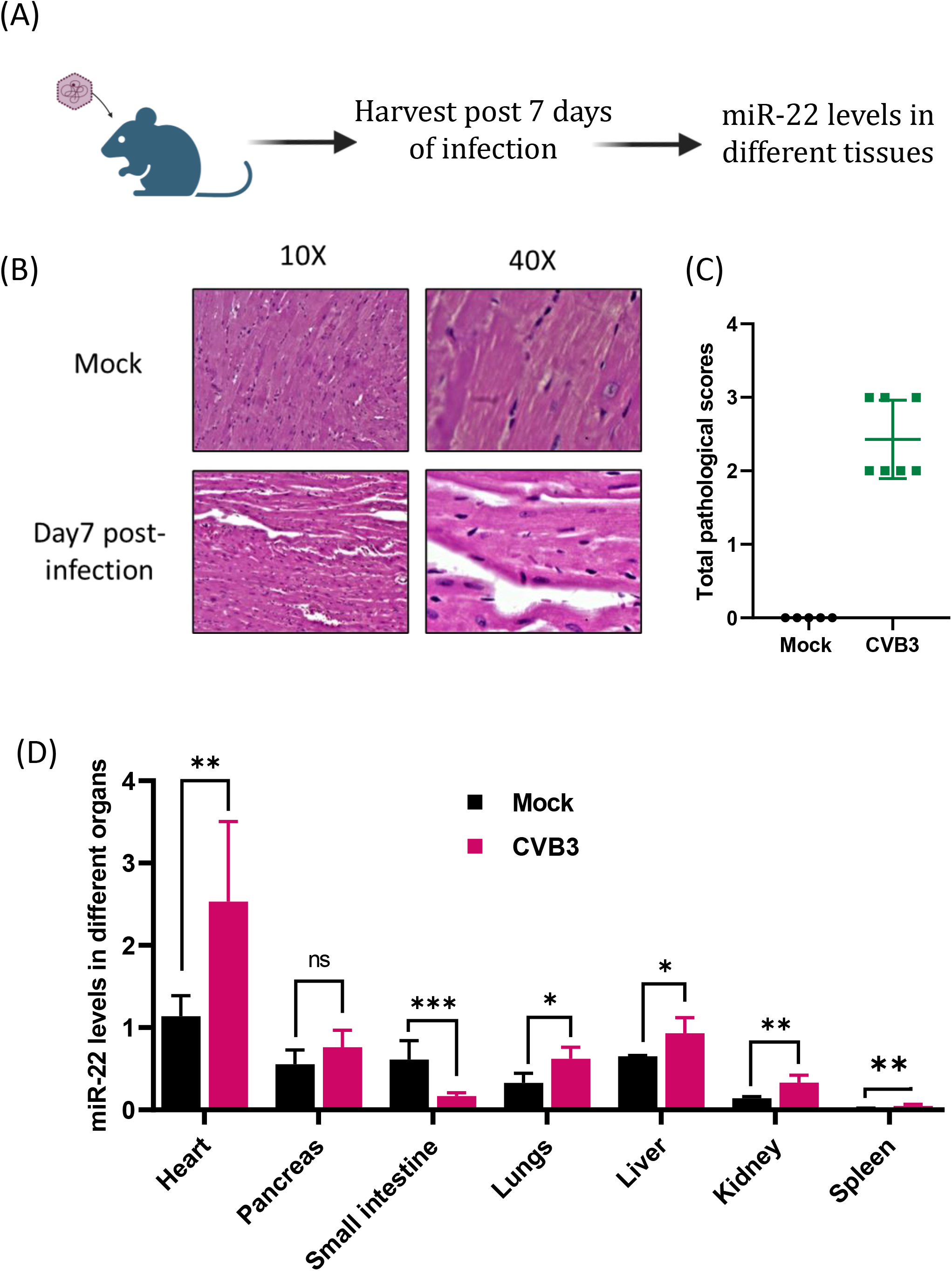
miR-22 levels determine the level of CVB3 infection in different organs. A) A schematic representation of the workflow for the consecutive experiment. 3-4 weeks old male BALB/c were either mock-infected or CVB3-infected (104 pfu). The animals were sacrificed after 7 days of infection and different organs were harvested. Total RNA was isolated from the different tissues and miR-22 levels were quantified using qPCR. B) Histopathology of Mock and CVB3-infected mice hearts. The CVB3-infected hearts showed clear sign of cardiac damage indicated by enlarged nuclei and fibrosis. C) Pathological score of the mock and CVB3-infected mice heart cross-sections. D) The bar graph represents the level of miR-22 in different organs of Mock and CVB3-infectd mice with respect to Mock-infected mice heart. Error bars represent standard deviation in five biological replicates. ****=p value<0.0001 ***=p value<0.001, **=p value<0.01, *=p value<0.05.

## DISCUSSION

Non-coding RNAs, such as miRNAs have proven to be essential host factors during many pathological conditions. In case of viral infections also, they can have either pro-viral or anti-viral role. During CVB3 infection, many miRNAs have been of importance in different phases, and they can affect the infection either by directly targeting the viral mRNA or targeting the host genes which might impact the infection. Only two reports are there for host miRNAs (miR-10a* and miR-342-5p) directly binding to CVB3 RNA and the binding the ORF region. In our study, we have delineated the role of another miRNA, miR-22, that binds to the CVB3 5’UTR and plays a crucial role in CVB3 RNA translation and also indirectly by targeting the host mRNA PCDH1.

We found that miR-22 binding to the CVB3 5’UTR is a mechanism for inhibiting CVB3 RNA translation. To be precise, when CVB3 releases its genomic RNA into the cell, several host proteins act as ITAFs and bind to the viral RNA in order to facilitate its translation. PSF, PTB and La proteins, which also share their binding site in the CVB3 RNA 5’UTR with miR-22, are known for promoting the translation of CVB3 RNA. One of which, La, also helps in the recruitment of the ribosome to the IRES. So, their binding promotes the CVB3 RNA translation, and it leads to the synthesis of viral proteins. After 4 hours of infection, when viral proteins, especially 2A protease, start to accumulate in the host cell, it triggers the induction of miR-22 and gives it a higher prospect of binding to the viral RNA. miR-22 binding to the 5’UTR displaces the ITAFs and inhibits the translation. Moreover, this increased level of miR-22 also affects its host target genes. We found that the increased level of miR-22 leads to reduction of Protocadherin-1 (PCDH1) which seems to be a pro-viral factor. So, a question arises why CVB3 will induce the levels of an antiviral miRNA? We believe that the levels of miR-22 are enhanced by CVB3 for its own benefit. The genomic RNA strand used for translation also has to be utilized for replication. And if the translation promoting factors keep occupying the CVB3 RNA, it will be difficult to make a switch from translation to replication. So, CVB3 increases miR-22 to aid in making the switch. However, miR-22 takes advantage of the situation and acts as an antiviral factor inhibiting pro-viral PCDH1, a repercussion the virus has to endure and overcome. We have been trying to solve this puzzle of regulation of different host factors in a time-dependent manner, which helps the virus in establishing a proper infection cycle. miR-22 contributes a small part in that puzzle (Figure 9A and B).

**Figure 9.**
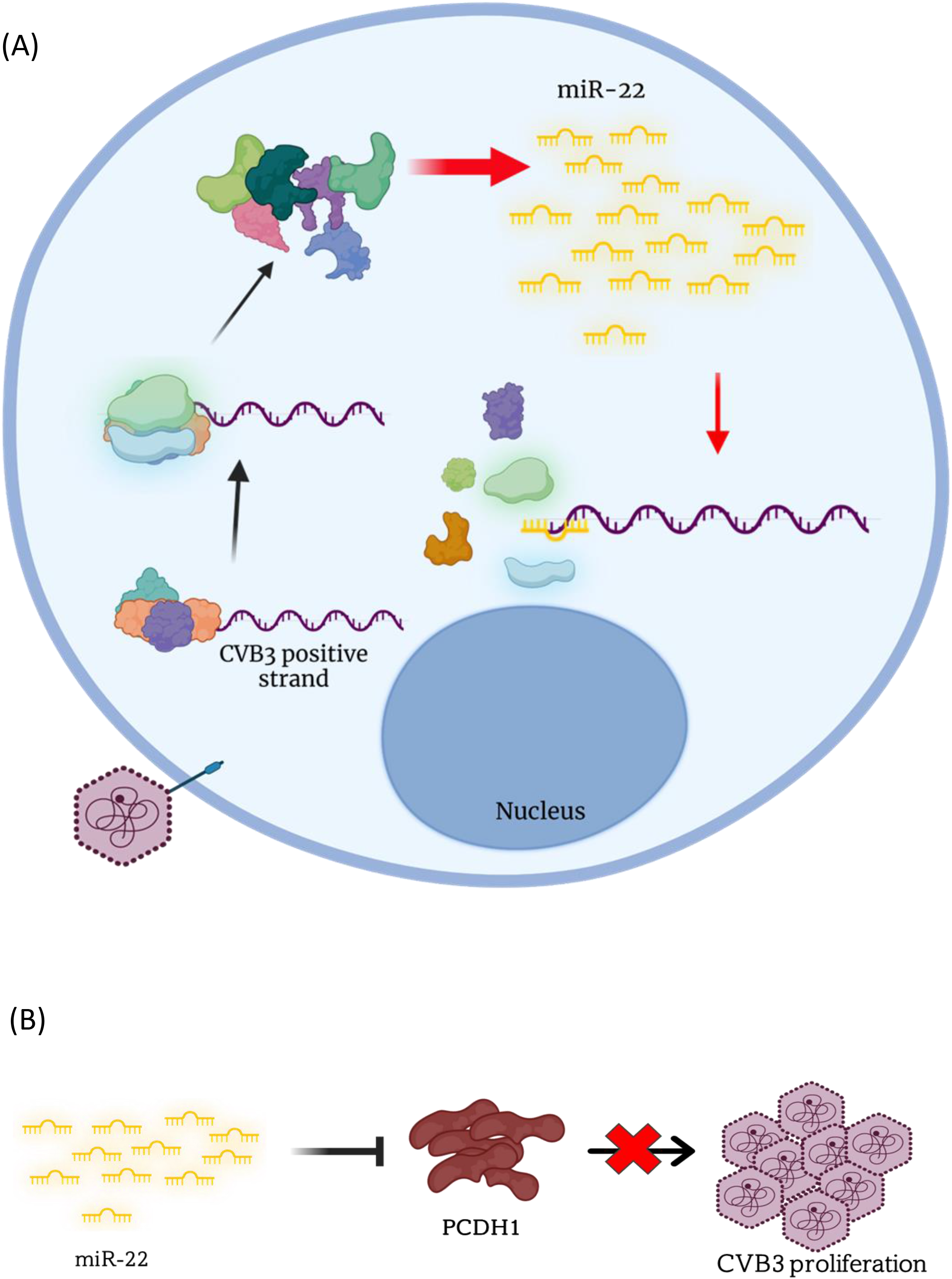
Models. A) CVB3 RNA translation inhibition by miR-22. CVB3 RNA released into the host cell cytoplasm is bound by different ITAFs such as La, PSF and PTB that promote its translation by recruiting ribosome. It leads to the production of CVB3 proteins. The accumulation of viral 2A protease in the cytoplasm acts as a trigger for enhancing the levels of miR-22, which now displaces the ITAFs bound to the CVB3 5’UTR, inhibiting its translation. B) The indirect effect of miR-22 during CVB3 infection by targeting PCDH1. During CVB3 infection, PCDH1 acts as a pro-viral host factor. However, the increased level of miR-22 upon CVB3 infection downregulates PCDH1 and inhibits its function.

## MATERIALS AND METHODS

### Cell lines and transfections

Hela cells were maintained in DMEM (Sigma) supplemented with 10% Foetal Bovine Serum (GIBCO, Invitrogen) and antibiotics. For transfection, 50 or 90% confluent monolayer of cells (depending on the experiment) was transfected with various plasmid constructs or RNAs using Lipofectamine 2000 (Invitrogen) or Turbofectamine in Opti-MEM (Invitrogen). Following a 5-hour incubation period, the culture medium was replaced with DMEM containing antibiotics and 10% FBS. Subsequently, at the designated time points, the cells were harvested and processed according to the specific requirements of the experiments.

### Plasmids and RNAs

pRib-CB3/T7-LUC construct was used for the preparation of CVB3 sub-genomic Replicon RNA. The construct linearized with SalI enzyme was used for in vitro transcription reactions using T7 polymerase as per manufacturer’s protocol.

pRib-CB3/T7 construct was used for the preparation of CVB3 infectious virus (Nancy strain). The construct linearized with MluI enzyme was used for in vitro transcription reactions using T7 polymerase as per manufacturer’s protocol.

pSUPER-miR22 and pSUPER-miR-21 constructs were used to overexpress miR-22 and miR-21, where the miRNA sequence was cloned between BglII and XhoI sites in pSUPER vector. For preparation of CVB3 5’UTR-FLuc, CVB3 5’UTR-FLuc-3’UTR, FLuc-CVB3 3’UTR RNA, respective sites were cloned in pcDNA3.1 vector. The clones were already available in the laboratory.

### Preparation of CVB3 infectious virus

The pRiB-T7-CB3 plasmid (A generous gift from Prof. Nora Chapman) containing CVB3 cDNA was used for infectious viral RNA preparation. This RNA was transfected into HeLa cells for CVB3 virus production. After 2-3 days of incubation, cells were lysed by three cycles of freeze-thawing and the cell suspension was collected in a 15 ml tube. Dead cells and cellular debris were removed by centrifugation at 5000 rpm for 10 minutes at 4℃. The supernatant containing the CVB3 virus was collected and aliquoted before being stored at -80℃ to maintain its infectivity. Freshly prepared virus was used to estimate plaque-forming units per millilitre using plaque assay in Vero cells.

### Virus infection in cell line

Cells were seeded at 90% confluency and before infection, washed with DMEM media without FBS. The required amount of virus is diluted in DMEM (without serum) and added to the cells. The dish is them swirled gently at regular intervals of 10 minutes for 45 min to 1 hour. This is the adsorption step. Next, remaining virus is removed, and fresh growth media is added to the cells.

### Virus infection in mice

3-4 weeks old male BALB/c mice were either injected with PBS for mock-infection or 10^4^ pfu of CVB3 intraperitoneally. The mice were sacrificed after 7 days of infection and the organs were harvested.

### Tagged RNA Affinity Pull down (TRAP) Assay

The method is based on adding MS2 RNA hairpin loops to a target RNA of interest, followed by co-expression of the MS2-tagged RNA together with the protein MS2 (which recognizes the MS2 RNA elements) fused to an affinity tag[32]. After purification of the MS2 RNP complex, the miRNAs present in the complex are identified. In this study, we have tagged the CVB3 5’UTR (WT or Mutant) with MS2 hairpins and have co-expressed it in HeLa cells along with the chimeric protein MS2-GST (glutathione S-transferase). After affinity purification using glutathione SH beads, the microRNAs present in the RNP complex were detected by qPCR.

### miR-22 Knockout cell line generation

pspCas9(BB)-2A-GFP(PX458) vector was used to clone miR-22 specific gRNAs. The clones were transfected in HeLa cell lines and 48h post transfection GFP-expressing cells were sorted in a 96-well plate by single cell sorting using FACS. The sorted cells were propagated and further miR-22 Knock out clones were screened and validated by sequencing and qPCR.

### MTT Assay

HeLa cells were transfected with pSUPER only, pSUPER-mir22, pSUPER-mir21 overexpression constructs or with Nsp Anti-miR, AntimiR-22 or AntimiR-21 and MTT [3-(4,5-dimethylthiazol-2-yl)-2,5-diphenyltetrazolium bromide] was added to the media 24 h post transfection. After 3 h of incubation, media was removed and cells were dissolved in DMSO, followed by measurement of absorbance at 550nm.

### In vitro transcriptions

RNA synthesis was performed in vitro using linearized constructs containing the T7 promoter. Two plasmids, pRib-CB3/T7-LUC, and pRIB-CB3/T7, were used, and they were linearized with SalI and MluI enzymes, respectively. The linearized DNA was then ethanol-precipitated and used as a template for RNA synthesis using T7 RNA polymerase (Fermentas). The transcription reactions were carried out under standard conditions (Fermentas protocol) using 1μg of linearized template DNA at 37℃ for 4hrs. After ethanol precipitation, RNAs were resuspended in nuclease free water.

### Luciferase assays

Transfected cells were harvested and lysed using 1X Passive Lysis Buffer (Promega). Cell lysates were centrifuged at 10,000 rpm for 10 minutes at 4℃. Supernatant was collected and used for taking the luciferase reading using Dual Luciferase kit (Promega) as per manufacturer’s protocol. Readings were normalized to total protein.

### Western Blot Analysis

The protein concentration from cell lysates was quantified using Bradford method. Equal amounts of protein were separated on 12% denaturing polyacrylamide gel electrophoresis (PAGE) gels and transferred to a nitrocellulose membrane (Pall Biosciences). The analysis of these proteins was conducted using specific antibodies.

### RNA immuno-precipitation

For RNA immune-precipitation, cells were transfected with CVB3 replicon RNA and 8 hours post-transfection, lysed with IP buffer containing 100 mM KCl, 5 mM MgCl2, 10 mM HEPES (pH 7.0), 0.5% NP-40, 1 mM DTT, and 100 U/ml RNase inhibitor. An equal amount of lysate was then incubated for 4 hours with protein G beads that had been pre-saturated with specific antibodies to form RNP complexes. Subsequently, the RNP complexes were washed thrice with the IP buffer, and the beads were resuspended in the same buffer. A 10% aliquot of this resuspended mixture was used for the analysis of proteins through western blotting, while the remaining portion was treated with 30 μg of proteinase K in the presence of 0.1% SDS at 50 °C for 30 minutes. Following this treatment, Trizol was added to these reactions for RNA isolation. The isolated RNA was then utilized for qPCR.

### RNA isolation and qPCR

Total RNA was extracted from cells at various time points using TRI Reagent (Sigma) following the standard protocol. After trypsin digestion, the cells were collected using PBS, and 250 μl of TRI Reagent was added to the cell pellet for proper lysis. Subsequently, 50 μl of chloroform was introduced to the tubes containing the lysed cells in TRI Reagent. After vigorous vertexing for 30 seconds, the tubes were centrifuged at 12,000 rpm for 20 minutes. The resulting supernatant was collected, and RNA precipitation was carried out by adding an equal volume of isopropanol. The RNA pellet was then washed with 70% ethanol (or 80% ethanol for miRNA quantification). Following this, DNaseI digestion was performed at 37°C for 30-45 minutes. Next, an equal volume of phenol-chloroform (1:1 ratio) was added to the solution, vortexed, and centrifuged at 10,000 rpm for 10 minutes. To the resulting supernatant, absolute ethanol (2.5 volumes), sodium acetate pH 5.2 (1/10th volume), and glycogen were added, and the mixture was kept at -80°C overnight for precipitation. After the overnight precipitation, the pellet was washed with 70% ethanol (or 80% ethanol for miRNA experiments), briefly air-dried, and then resuspended in DEPC-treated water.

### Polysome profiling

The wild type or miR22 knockout cells HeLa were cultured in 100-mm dishes and infected with CVB3 virus. After 8 hours of incubation, the cells were treated with cycloheximide (100 μg/ml) for 10 minutes at 37°C. Following this, the cells were washed with ice-cold PBS containing cycloheximide and then with hypotonic buffer [5 mM Tris-HCl (pH 7.5), 1.5 mM KCl, 5 mM MgCl2, and 100 μg/ml cycloheximide]. The cells were then scraped in ice-cold lysis buffer [5 mM Tris-HCl (pH 7.5), 1.5 mM KCl, 5 mM MgCl2, 100 μg/ml cycloheximide, 1 mM DTT, 200 U/ml RNasin, 200 μg/ml tRNA, 0.5% Triton X-100, 0.5% sodium deoxycholate, and 1x protease inhibitor cocktail] and kept on ice for 15 minutes. After 15 minutes, the lysate’s KCl concentration was adjusted to a final concentration of 150 mM. The cell lysate was then centrifuged at 3,000 x g for 8 minutes at 4°C, and the resulting supernatant was collected and either processed immediately or flash frozen and stored at -80°C for later use. Subsequently, 1-1.5 μg of total protein was loaded onto a 10% to 50% sucrose gradient and centrifuged at 36,000 rpm for 2 hours at 4°C using an SW41 rotor (Beckman). Polysome profiles were visualized, and the fractions were collected using a Polysome profiler (BioComp). Individual fractions were then used for RNA isolation and qPCR.

## AUTHOR CONTRIBUTIONS

PR, BG, and SD: Conception and design of studies, analysis, and interpretation. PR and SD: article writing. PR, BG, SV, SB, RS, and AP for performing the experiments.

## ACKNOWLEDGEMENTS

We thank Frank van Kuppeveld and Nora Chapman for various constructs. We thank the Centre for Infectious Disease Research, Indian Institute of Science (IISc), for allowing use of the biosafety level 3 (BSL3) facility and the Biological Science department, IISc, for use of the FACS facility. We also thank Deepak Saini, Department of Molecular Reproduction, Development and Genetics, IISc, for providing the HeLa cell line, Umesh Varshney, Department of Microbiology and Cell Biology, for allowing the usage of laboratory facilities for polysome analysis.

P.R., S.V., and A.P. is supported by the Ministry of Human Resource Development (MHRD). Funding was provided by a D.S. Kothari fellowship from University Grants Commission, Government of India to B.G. Financial help from DBT-IISc Partnership Program is also acknowledged. S.D. acknowledges the J C Bose fellowship from Department of Science and Technology, Government of India. The work was also supported by a research grant from Department of Biotechnology, Government of India.

## DATA AVAILABILITY STATEMENT

Data available within the article or its supplementary materials.

## REFERENCES

1. Harris, K. G.; Coyne, C. B., Death waits for no man--does it wait for a virus? How enteroviruses induce and control cell death. Cytokine Growth Factor Rev 2014, 25, (5), 587–96.

2. Garmaroudi, F. S.; Marchant, D.; Hendry, R.; Luo, H.; Yang, D.; Ye, X.; Shi, J.; McManus, B. M., Coxsackievirus B3 replication and pathogenesis. Future Microbiol 2015, 10, (4), 629–53.

3. Alirezaei, M.; Flynn, C. T.; Wood, M. R.; Whitton, J. L., Pancreatic acinar cell-specific autophagy disruption reduces coxsackievirus replication and pathogenesis in vivo. Cell Host Microbe 2012, 11, (3), 298–305.

4. Fairweather, D.; Stafford, K. A.; Sung, Y. K., Update on coxsackievirus B3 myocarditis. Current opinion in rheumatology 2012, 24, (4), 401–7.

5. Zhang, C.; Xiong, Y.; Zeng, L.; Peng, Z.; Liu, Z.; Zhan, H.; Yang, Z., The Role of Non-coding RNAs in Viral Myocarditis. 2020, 10.

6. Hemida, M. G.; Ye, X.; Zhang, H. M.; Hanson, P. J.; Liu, Z.; McManus, B. M.; Yang, D., MicroRNA-203 enhances Coxsackievirus B3 replication through targeting zinc finger protein-148. Cellular and Molecular Life Sciences 2013, 70, (2), 277–291.

7. Germano, J. F.; Sawaged, S.; Saadaeijahromi, H.; Andres, A. M.; Feuer, R.; Gottlieb, R. A.; Sin, J., Coxsackievirus B infection induces the extracellular release of miR-590-5p, a proviral microRNA. Virology 2019, 529, 169–176.

8. Ye, X.; Hemida, M. G.; Qiu, Y.; Hanson, P. J.; Zhang, H. M.; Yang, D., MiR-126 promotes coxsackievirus replication by mediating cross-talk of ERK1/2 and Wnt/β-catenin signal pathways. Cellular and molecular life sciences: CMLS 2013, 70, (23), 4631–44.

9. Corsten, M. F.; Heggermont, W.; Papageorgiou, A.-P.; Deckx, S.; Tijsma, A.; Verhesen, W.; van Leeuwen, R.; Carai, P.; Thibaut, H.-J.; Custers, K.; Summer, G.; Hazebroek, M.; Verheyen, F.; Neyts, J.; Schroen, B.; Heymans, S., The microRNA-221/-222 cluster balances the antiviral and inflammatory response in viral myocarditis. European Heart Journal 2015, 36, (42), 2909–2919.

10. Jiang, D.; Li, M.; Yu, Y.; Shi, H.; Chen, R., microRNA-34a aggravates coxsackievirus B3-induced apoptosis of cardiomyocytes through the SIRT1-p53 pathway. J Med Virol 2019, 91, (9), 1643–1651.

11. Zhang, X.; Gao, X.; Hu, J.; Xie, Y.; Zuo, Y.; Xu, H.; Zhu, S., ADAR1p150 Forms a Complex with Dicer to Promote miRNA-222 Activity and Regulate PTEN Expression in CVB3-Induced Viral Myocarditis. International journal of molecular sciences 2019, 20, (2).

12. Zhang, B. Y.; Zhao, Z.; Jin, Z., Expression of miR-98 in myocarditis and its influence on transcription of the FAS/FASL gene pair. Genetics and molecular research: GMR 2016, 15, (2).

13. He, F.; Xiao, Z.; Yao, H.; Li, S.; Feng, M.; Wang, W.; Liu, Z.; Liu, Z.; Wu, J., The protective role of microRNA-21 against coxsackievirus B3 infection through targeting the MAP2K3/P38 MAPK signaling pathway. Journal of translational medicine 2019, 17, (1), 335.

14. He, J.; Yue, Y.; Dong, C.; Xiong, S., MiR-21 confers resistance against CVB3-induced myocarditis by inhibiting PDCD4-mediated apoptosis. Clinical and investigative medicine. Medecine clinique et experimentale 2013, 36, (2), E103–11.

15. Wang, L.; Qin, Y.; Tong, L.; Wu, S.; Wang, Q.; Jiao, Q.; Guo, Z.; Lin, L.; Wang, R.; Zhao, W.; Zhong, Z., MiR-342-5p suppresses coxsackievirus B3 biosynthesis by targeting the 2C-coding region. Antiviral research 2012, 93, (2), 270–279.

16. Tong, L.; Lin, L.; Wu, S.; Guo, Z.; Wang, T.; Qin, Y.; Wang, R.; Zhong, X.; Wu, X.; Wang, Y.; Luan, T.; Wang, Q.; Li, Y.; Chen, X.; Zhang, F.; Zhao, W.; Zhong, Z., MiR-10a* up-regulates coxsackievirus B3 biosynthesis by targeting the 3D-coding sequence. Nucleic acids research 2013, 41, (6), 3760–3771.

17. Ray, P. S.; Das, S., La autoantigen is required for the internal ribosome entry site-mediated translation of Coxsackievirus B3 RNA. Nucleic acids research 2002, 30, (20), 4500–4508.

18. Dave, P.; George, B.; Sharma, D. K.; Das, S., Polypyrimidine tract-binding protein (PTB) and PTB-associated splicing factor in CVB3 infection: an ITAF for an ITAF. Nucleic Acids Research 2017, 45, (15), 9068–9084.

19. Verma, B.; Ponnuswamy, A.; Gnanasundram, S. V.; Das, S., Cryptic AUG is important for 48S ribosomal assembly during internal initiation of translation of coxsackievirus B3 RNA. Journal of General Virology 2011, 92, (10), 2310–2319.

20. Xu, D.; Takeshita, F.; Hino, Y.; Fukunaga, S.; Kudo, Y.; Tamaki, A.; Matsunaga, J.; Takahashi, R. U.; Takata, T.; Shimamoto, A.; Ochiya, T.; Tahara, H., miR-22 represses cancer progression by inducing cellular senescence. The Journal of cell biology 2011, 193, (2), 409–24.

21. Zhang, X.; Li, Y.; Wang, D.; Wei, X., miR-22 suppresses tumorigenesis and improves radiosensitivity of breast cancer cells by targeting Sirt1. Biological research 2017, 50, (1), 27.

22. Fan, T.; Wang, C. Q.; Li, X. T.; Yang, H.; Zhou, J.; Song, Y. J., MiR-22-3p Suppresses Cell Migration and Invasion by Targeting PLAGL2 in Breast Cancer. Journal of the College of Physicians and Surgeons--Pakistan: JCPSP 2021, 31, (8), 937–940.

23. Wu, H.; Liu, J.; Zhang, Y.; Li, Q.; Wang, Q.; Gu, Z., miR-22 suppresses cell viability and EMT of ovarian cancer cells via NLRP3 and inhibits PI3K/AKT signaling pathway. Clinical & translational oncology: official publication of the Federation of Spanish Oncology Societies and of the National Cancer Institute of Mexico 2021, 23, (2), 257–264.

24. Yang, Q. Y.; Yang, K. P.; Li, Z. Z., MiR-22 restrains proliferation of rheumatoid arthritis by targeting IL6R and may be concerned with the suppression of NF-κB pathway. The Kaohsiung journal of medical sciences 2020, 36, (1), 20–26.

25. Xia, P.; Chen, J.; Liu, Y.; Cui, X.; Wang, C.; Zong, S.; Wang, L.; Lu, Z., MicroRNA-22-3p ameliorates Alzheimer’s disease by targeting SOX9 through the NF-κB signaling pathway in the hippocampus. Journal of neuroinflammation 2022, 19, (1), 180.

26. Ji, Q.; Wang, X.; Cai, J.; Du, X.; Sun, H.; Zhang, N., MiR-22-3p Regulates Amyloid β Deposit in Mice Model of Alzheimer’s Disease by Targeting Mitogen-activated Protein Kinase 14. Current neurovascular research 2019, 16, (5), 473–480.

27. Huang, Z.-P.; Wang, D.-Z., miR-22 in cardiac remodeling and disease. Trends Cardiovasc Med 2014, 24, (7), 267–272.

28. Huang, Z.-P.; Chen, J.; Seok, H. Y.; Zhang, Z.; Kataoka, M.; Hu, X.; Wang, D.-Z., MicroRNA-22 regulates cardiac hypertrophy and remodeling in response to stress. Circ Res 2013, 112, (9), 1234–1243.

29. George, B.; Dave, P.; Rani, P.; Behera, P.; Das, S., Cellular Protein HuR Regulates the Switching of Genomic RNA Templates for Differential Functions during the Coxsackievirus B3 Life Cycle. J Virol 2021, 95, (21), e0091521.

30. Ye, Z.; Yang, Y.; Wei, Y.; Li, L.; Wang, X.; Zhang, J., PCDH1 promotes progression of pancreatic ductal adenocarcinoma via activation of NF-κB signalling by interacting with KPNB1. Cell death & disease 2022, 13, (7), 633.

31. Jangra, R. K.; Herbert, A. S.; Li, R.; Jae, L. T.; Kleinfelter, L. M.; Slough, M. M.; Barker, S. L.; Guardado-Calvo, P.; Román-Sosa, G.; Dieterle, M. E.; Kuehne, A. I.; Muena, N. A.; Wirchnianski, A. S.; Nyakatura, E. K.; Fels, J. M.; Ng, M.; Mittler, E.; Pan, J.; Bharrhan, S.; Wec, A. Z.; Lai, J. R.; Sidhu, S. S.; Tischler, N. D.; Rey, F. A.; Moffat, J.; Brummelkamp, T. R.; Wang, Z.; Dye, J. M.; Chandran, K., Protocadherin-1 is essential for cell entry by New World hantaviruses. Nature 2018, 563, (7732), 559–563.

32. Yoon, J. H.; Srikantan, S.; Gorospe, M., MS2-TRAP (MS2-tagged RNA affinity purification): tagging RNA to identify associated miRNAs. *Methods (San Diego*, Calif.) 2012, 58, (2), 81–7.

