## Supplemental Figure 1 for "microRNA-22 displaces ITAFs from the 5’UTR and inhibit the translation of Coxsackievirus B3 RNA"

### Slide 1
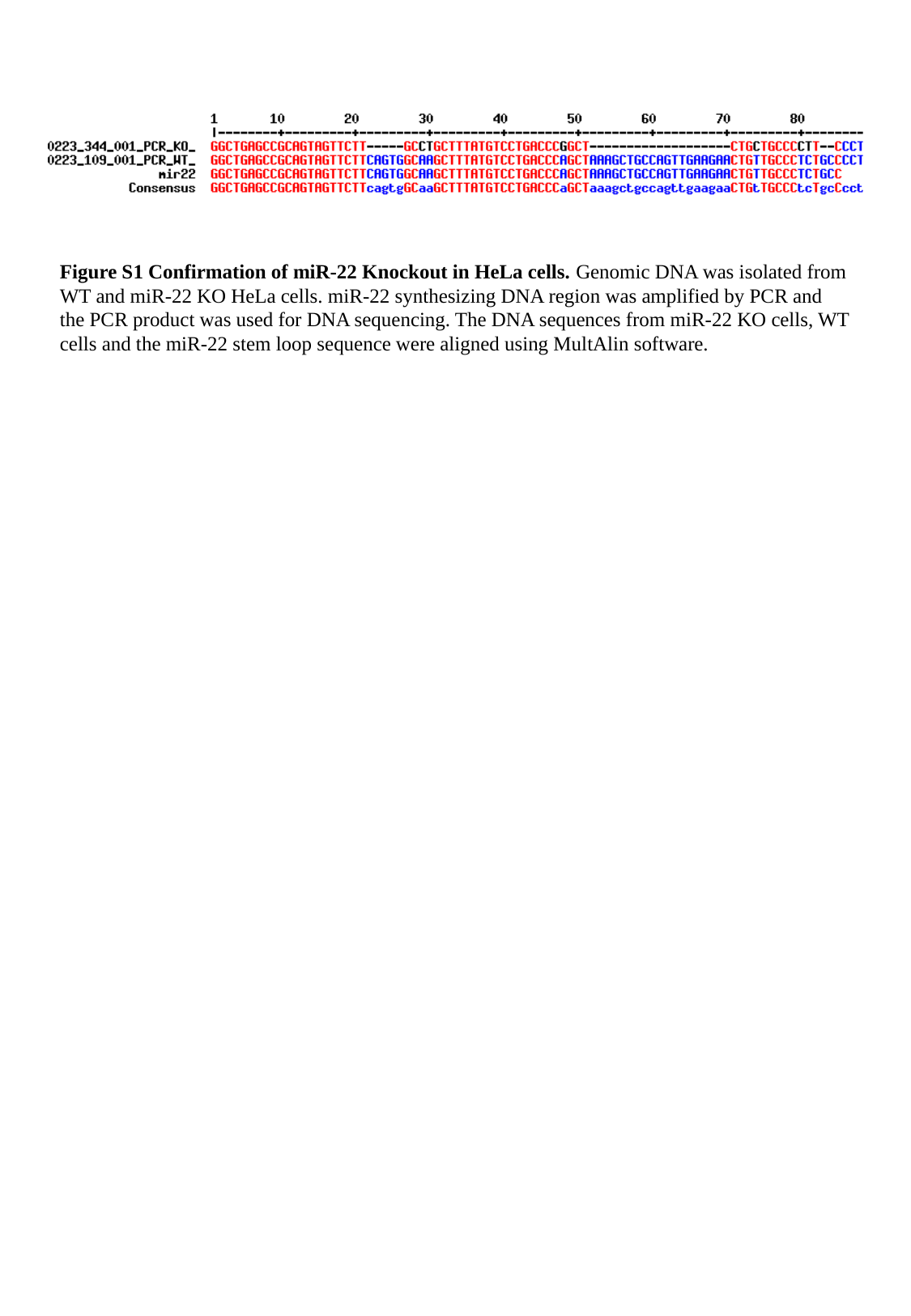

Figure S1 Confirmation of miR-22 Knockout in HeLa cells. Genomic DNA was isolated from WT and miR-22 KO HeLa cells. miR-22 synthesizing DNA region was amplified by PCR and the PCR product was used for DNA sequencing. The DNA sequences from miR-22 KO cells, WT cells and the miR-22 stem loop sequence were aligned using MultAlin software.
