## Supplementary figures and images for "microRNA-22 displaces ITAFs from the 5’UTR and inhibit the translation of Coxsackievirus B3 RNA"

### Supplemental Figure 2

#### Slide 1
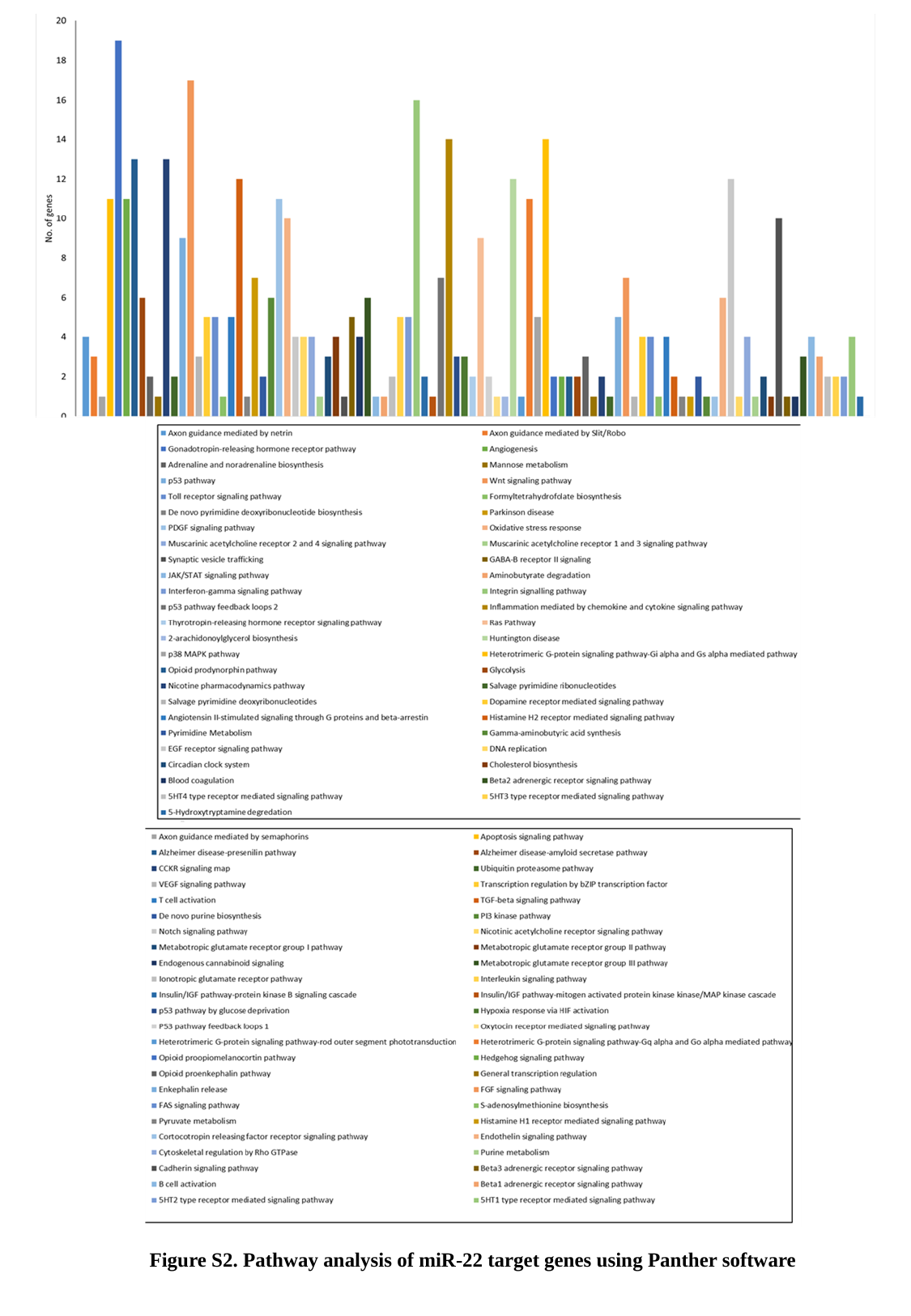

Figure S2. Pathway analysis of miR-22 target genes using Panther software
