## Supplemental table 1 for "microRNA-22 displaces ITAFs from the 5’UTR and inhibit the translation of Coxsackievirus B3 RNA"

### Slide 1
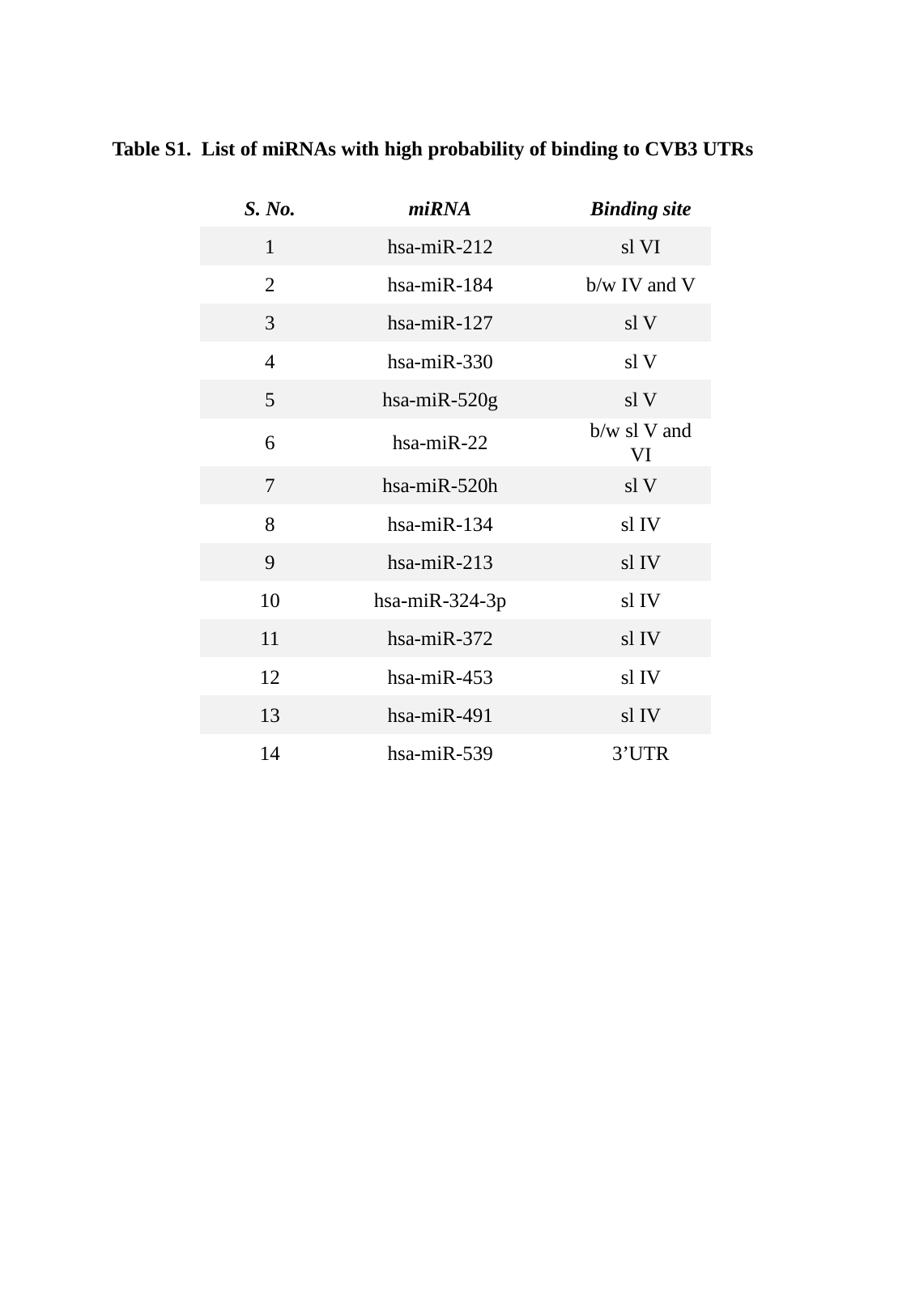

Table S1. List of miRNAs with high probability of binding to CVB3 UTRs
| S. No. | miRNA | Binding site |
| --- | --- | --- |
| 1 | hsa-miR-212 | sl VI |
| 2 | hsa-miR-184 | b/w IV and V |
| 3 | hsa-miR-127 | sl V |
| 4 | hsa-miR-330 | sl V |
| 5 | hsa-miR-520g | sl V |
| 6 | hsa-miR-22 | b/w sl V and VI |
| 7 | hsa-miR-520h | sl V |
| 8 | hsa-miR-134 | sl IV |
| 9 | hsa-miR-213 | sl IV |
| 10 | hsa-miR-324-3p | sl IV |
| 11 | hsa-miR-372 | sl IV |
| 12 | hsa-miR-453 | sl IV |
| 13 | hsa-miR-491 | sl IV |
| 14 | hsa-miR-539 | 3’UTR |
