## Supplemental table 2 for "microRNA-22 displaces ITAFs from the 5’UTR and inhibit the translation of Coxsackievirus B3 RNA"

### Slide 1
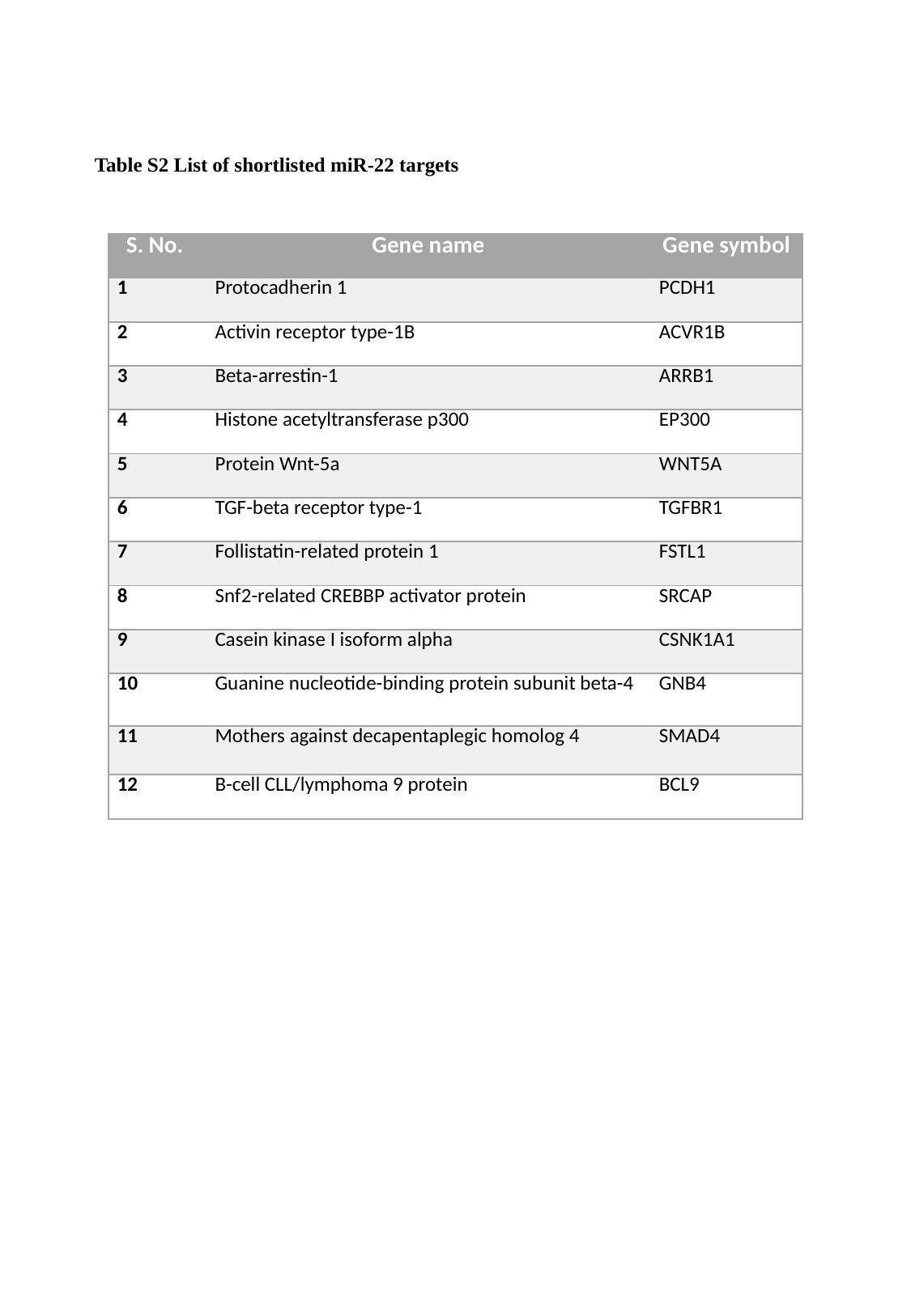

Table S2 List of shortlisted miR-22 targets
| S. No. | Gene name | Gene symbol |
| --- | --- | --- |
| 1 | Protocadherin 1 | PCDH1 |
| 2 | Activin receptor type-1B | ACVR1B |
| 3 | Beta-arrestin-1 | ARRB1 |
| 4 | Histone acetyltransferase p300 | EP300 |
| 5 | Protein Wnt-5a | WNT5A |
| 6 | TGF-beta receptor type-1 | TGFBR1 |
| 7 | Follistatin-related protein 1 | FSTL1 |
| 8 | Snf2-related CREBBP activator protein | SRCAP |
| 9 | Casein kinase I isoform alpha | CSNK1A1 |
| 10 | Guanine nucleotide-binding protein subunit beta-4 | GNB4 |
| 11 | Mothers against decapentaplegic homolog 4 | SMAD4 |
| 12 | B-cell CLL/lymphoma 9 protein | BCL9 |
